# The language oscillogenome

**DOI:** 10.1101/114033

**Authors:** Elliot Murphy, Antonio Benítez-Burraco

## Abstract

Language has been argued to arise, both ontogenetically and phylogenetically, from specific patterns of brain wiring. We argue that it can further be shown that core features of language processing emerge from particular phasal and cross-frequency coupling properties of neural oscillations; what has been referred to as the language ‘oscillome’. It is expected that basic aspects of the language oscillome result from genetic guidance, what we will here call the language ‘oscillogenome’, for which we will put forward a list of candidate genes. We have considered genes for altered brain rhythmicity in conditions involving language deficits (autism spectrum disorders, schizophrenia, specific language impairment and dyslexia) for which we have confident genome-oscillome-phenome connections. These selected genes map on to aspects of brain function, particularly on to neurotransmitter function. Our aim is to propose a set of biologically robust genome-to-language linking hypotheses that, given testing, would grant causal and explanatory power to brain rhythms with respect to language processing.

## 1. Introduction

Which genes regulate mental processes? This is surely one of the most pivotal questions in contemporary neurobiology. In their Foreword to a recent volume on birdsong and biolinguistics, Berwick and Chomsky (2015: x) discuss the potential for one particular gene, *FoxP2*, to contribute to debates about the evolution of our most complex mental capacity, language, commenting that ‘[h]ow far one can drive this genomic work upward into neuronal assemblies – ultimately, the dissection of the underlying circuitry responsible for vocal production – remains to be seen’.

The present paper will serve as the next step in this biolinguistic approach to language, documenting the genes implicated in oscillatory activity during language processing as a means of establishing causal genome-oscillome linking hypotheses. As is standard in candidate-gene approach studies, we have examined cognitive conditions entailing both language deficits and oscillatory anomalies as a way to identify promising candidates. We have focused on schizophrenia (SZ) and autism spectrum disorders (ASD), which entail language impairment mostly at the syntax-semantics interface, and also on specific language impairment (SLI) and developmental dyslexia (DD), which entail language impairment mostly with prosody and phonology. For a proposal of how to explain these deficits in oscillatory terms, see Benítez-Burraco and Murphy (2016), Murphy and Benítez-Burraco (2016a, b), and Jiménez-Bravo et al. (2017). At the core of these proposals is the assumption that particular computational and representational properties can be attributed to neural oscillations. This continues a recent line of research which has drawn the following conclusions: Computational operations of language can be decomposed into generic processes (Murphy 2015a); these generic processes interact in dynamic ways and can be implemented via neural oscillations (Murphy 2015b); these oscillations implement a multiplexing algorithm for the combination and interpretation of linguistic representations (Murphy 2016a); this multiplexing algorithm appears to be species-specific (Murphy 2016b). The long-standing conclusions concerning the species-specificity of language therefore come full circle through a human-specific oscillatory code. What we have argued is that, in essence, although most of the nerve tracks and regions which differ in these pathological conditions are implicated in language processing, neural oscillations provide a more reliable explanatory level of the language deficits exhibited by the affected populations. Moving beyond this now requires an examination of the genes responsible for the brain’s oscillatory behavior.

The genetic basis of neural oscillations more broadly likely stems from regulatory genes controlling the brain’s neurochemistry (Begleiterand Porjesz 2006). Oscillations represent highly heritable traits (van Beijsterveldt et al. 1996, Linkenkaer-Hansen et al. 2007, Hall et al. 2011) that are an interesting combination of being less complex but more proximal to gene function than standard cognitive or diagnostic labels. In what follows, we first provide a functional characterization of candidate genes for the language oscillogenome, with a focus on their biological significance and functions. We then discuss the contribution of these genes to language processing, and sketch genome-to-oscillome-to-language links. With this aim, we will consider the brain areas in which they are expressed, the brain rhythms they have been related to, and the role of these areas and rhythms in language processing. Our goal is to understand how these genes contribute to language processing and how mutations in these genes result in language impairments, with a focus on normal or abnormal oscillatory activity. We conclude with a brief discussion concerning future perspectives for finding links between genes, brain rhythms, and language.

## 2. A draft of the language oscillogenome

In order to achieve our objective of drafting the language oscillogenome, we first gathered via systematic literature review and database searches a list of potential candidates. We selected genes that i) are associated with language disorders (DD and SLI) or to language dysfunction in cognitive disorders entailing language deficits (SZ and ASD), and ii) are known to play a role in brain rhythmicity and/or are candidates for conditions entailing brain dysrhythmias, like epilepsy. As noted in the introduction, we have chosen these four clinical conditions because of three main reasons. First, in our previous work (Benítez-Burraco and Murphy 2016, Murphy and Benítez-Burraco 2016a,b, Jiménez-Bravo et al. 2017), we have provided characterizations of their linguistic profile in terms of an abnormal suite of brain rhythms and we have advanced some promising gene-oscillations-language links. Second, language impairment in these conditions relate to core aspects of language (and of language processing in the brain), in particular, to the interface between syntax and semantics, and between syntax and phonology. Third, we have already proposed a list of candidate genes for the oscillopathic profile of language dysfunction in these conditions.

For SZ we have mostly relied on the Schizophrenia Database (SZDB) (http://www.szdb.org). We have considered 679 candidates based on different source of evidence: candidates resulting from genome-wide association studies (GWAs), genes affected by copy number variant (CNV), genes identified by convergent functional genomics, and genes identified by linkage and association studies. Within these genes, we have identified those that have been found to play a role in language development (and potentially evolution), and to play as well some known role in brain rhythmicity, as discussed in Murphy and Benítez-Burraco (2016a) and Murphy and Benítez-Burraco (2016b). For ASD we have relied mostly on the SFARI database (https://sfari.org), which currently includes 881 genes related to the disorder, based on different levels of evidence (genes bearing rare single variants, disruptions/mutations, or small deletions/duplications; candidates resulting from genetic association studies, particularly, GWAs; genes resulting from functional approaches; and genes with CNV associated with ASD).Within these genes, we have equally focused on those highlighted in Benítez-Burraco and Murphy (2016) and Murphy and Benítez-Burraco (2016b) as important for language development and evolution. For DD, we have mostly relied on the last updated list of candidates for this condition, as provided by Paracchini et al (2016), which includes genes resulting from candidate association studies, GWAs, quantitative GWAs, CNV studies, and next-generation sequencing (NGS) analyses, although we have also surveyed the literature looking for additional candidates. As before, we selected among these genes those with a known role in brain oscillations. Finally, for SLI, we have mostly relied on the literature review provided by Chen et al. (2017) and on the literature survey and results provided by Pettigrew et al (2016), which contain candidates resulting from linkage analyses, GWA studies, and NGS analyses. As with DD, we surveyed the literature looking for other candidates for this condition. Among these genes, we selected, as noted, those with a stablished role in brain rhythmicity.

Our list of potential candidates for the language oscillogenome is shown in Table 1.

**Table 1.**
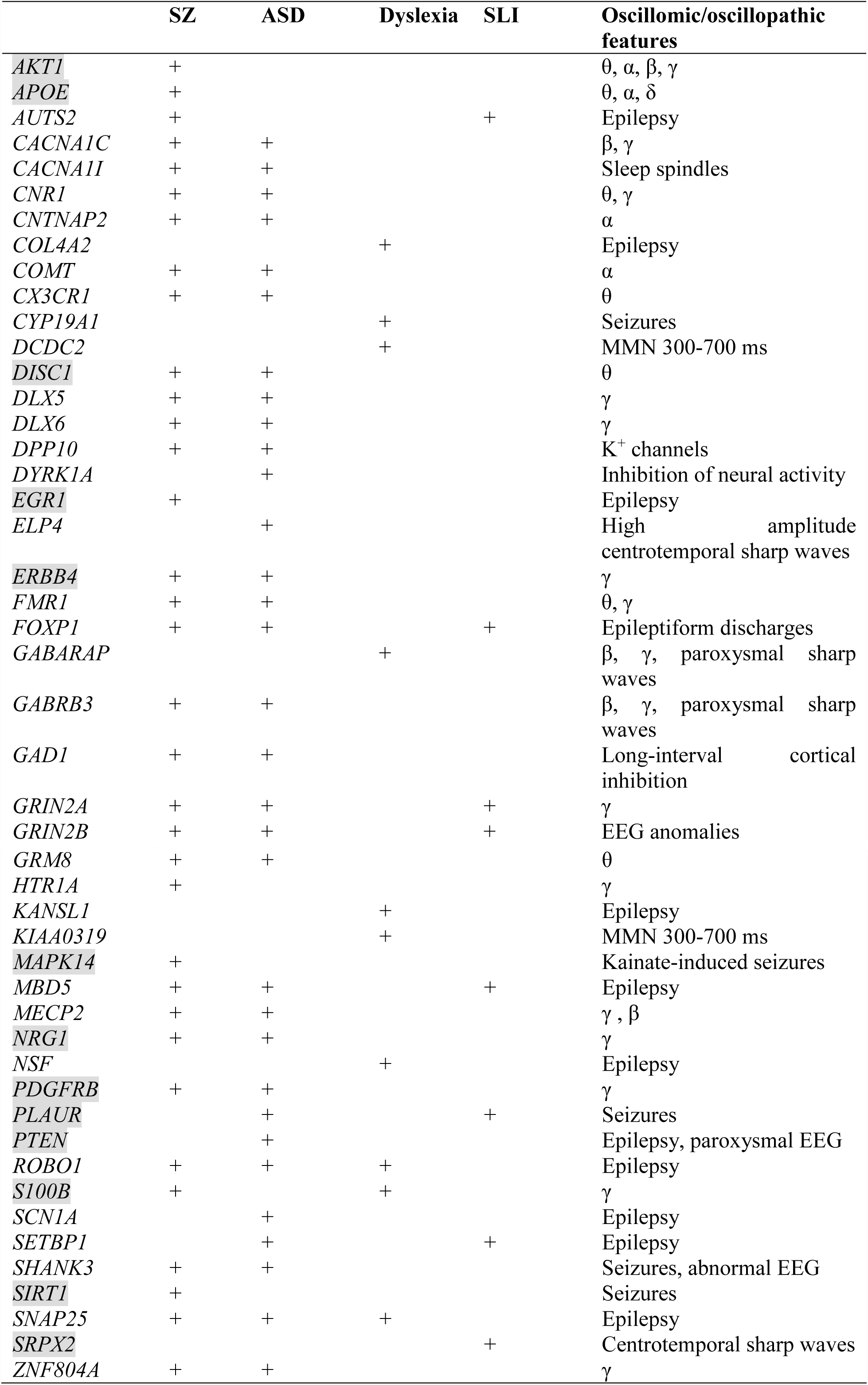
The set of candidates comprising the language oscillogenome. The genes shadowed in grey are those found strongly interconnected in our network, as shown in Figure 1 and discussed in the text.

We expected that the 48 genes we highlight here as part of the shared signature of abnormal brain oscillations associated with language deficits exhibit some kind of functional relationship, and map on to particular regulatory pathways, cell types or functions, or facets of brain development and function of relevance for language and the etiopathogenesis of language impairment in the clinical conditions we have mentioned. Accordingly, we used String 10.5 (www.string-db.org) for examining potential functional links among the proteins encoded by our candidates. String 10 is a predictive tool of direct/physical and indirect/functional associations between proteins that are derived from four sources: genomic context, high-throughput experiments, conserved coexpression, and the knowledge previously gained from text mining (Szklarczyk et al. 2015). In order to uncover potential clusters within our network, a MCL clustering algorithm (inflation parameter = 2) was applied to the distance matrix obtained from the String global scores. The MCL algorithm was used because it is remarkably robust to graph alterations and provides with better extractions of complexes from interaction networks (Brohée and van Helden 2006). As shown in Figure 1, our network comprises several clusters of interest. Also, several proteins (NRG1, ERBB4, PDGFRB, EGR1, APOE, AKT1, MAPK14, PTEN, DISC1, SIRT1, PLAUR, SRPX2, and S100B) are found strongly interconnected: they belong to the same cluster and the confidence values of most of the edges are high (0.700) or at the highest (0.900). Hence, we expect them to be key, core components of the regulatory network involved in different steps of neural development and function important for language processing, particularly in brain oscillatory activity.

**Figure 1.**
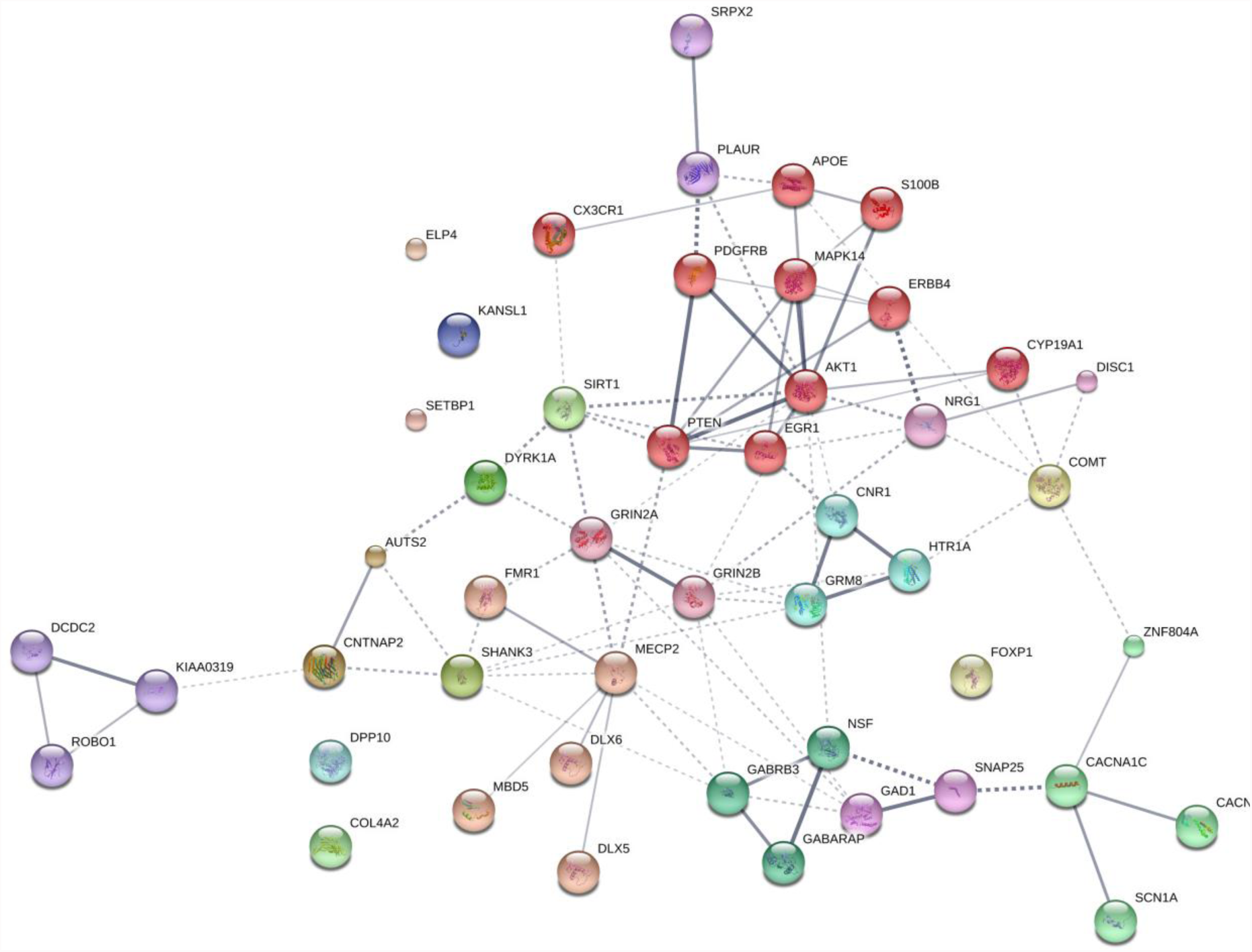
Protein interaction network. The diagram shows the network of known and predicted interactions among proteins encoded by genes proposed as candidates for the language oscillogenome (Table 1). The network was drawn with String (version 10.0; Szklarczyk et al., 2015) license-free software (<u>http://string-db.org/</u>), using the confidence visualization. It contains 48 nodes and 86 edges, with an average node degree of 3.58 and an average local clustering coefficient of 0.384. Colored nodes symbolize proteins included in the query (small nodes are for proteins with unknown 3D structure, while large nodes are for those with known structures). The line thickness of the edges indicates the strength of data support. The medium confidence value was .0400 (a 40% probability that a predicted link exists between two enzymes in the same metabolic map in the KEGG database: <u>http://www.genome.jp/kegg/pathway.html</u>). Each protein cluster is represented with a different color of the nodes. Inter-cluster edges are represented by dashed-lines. The diagram only represents the potential connectivity between the involved proteins, which has to be mapped onto particular biochemical networks, signaling pathways, cellular properties, aspects of neuronal function, or cell-types of interest (see the main text for details and Table 2 for a GO analysis).

NRG1 is a membrane glycoprotein that mediates cell-cell signaling and that contributes to the regulation of neural proliferation in the subventricular zone (Ghashghaei et al. 2006), thalamocortical axon pathfinding (López-Bendito et al. 2006), and glutamatergic and dopaminergic neurotransmission in the thalamus and the striatum (Newell et al. 2013). NRG1 and its receptor ERBB4 regulate as well how GABAergic interneurons migrate from ganglionic eminences to cortex (Li et al. 2012). Additionally, they play a key role in synchronizing neural oscillations in the cortex. Specifically, they enhance the synchrony of pyramidal neurons via presynaptic interneurons, increase the synchrony between pairs of fast-spiking interneurons and non-fast-spiking interneurons in prefrontal cortex, and enhance kainate-induced gamma oscillations in vivo (Hou et al. 2014). Risk alleles of *NRG1* have been found to correlate with semantic (but not to lexical) verbal fluency in SZ, and with a decreased activation in the right middle temporal gyri and the anterior cingulate gyrus, as well as the left inferior frontal of schizophrenic patients (Kircher et al. 2009), but also with a reduction of the left superior temporal gyrus volumes (Tosato et al. 2012). Risk polymorphisms of the gene are also implicated in enhanced memory, IQ scores and linguistic abilities in patients with bipolar disorder (Rolstad et al. 2015). Interestingly too, *Nrg1*(+/-) mice exhibit decreased social activity which mimic the social deficits observed in autistic patients (Ehrlichman et al., 2009).

Epistatic interactions between *Nrg1* and *Akt1* have been found to regulate aspects of behavioral phenotypes and social functions in genetic mouse models of SZ; specifically, double mutant mice exhibit impaired episodic-like memory and impaired sociability, as well as reduced ultrasonic vocalization calls (Huang et al 2015). Likewise, epistatic interactions between SZ-risk polymorphisms of *AKT1* and *COMT* (discussed below) are relevant for human medial temporal lobe structure and memory function (Tan et al. 2012a). Functional polymorphisms of *AKT1* have been related to the dopaminergic signalling in the prefrontal-striatal circuits responsible for the manipulative component of the working memory (Tan et al, 2012b). Specifically, an *AKT1* allele has been associated with verbal learning and memory (Pietiläinen et al. 2009). Genetic deletion of *Akt1* in mice impairs hippocampal long-term potentiation and affects spatial learning, suggesting that *AKT1* contributes to regulate hippocampal neuroplasticity and cognition (Balu et al. 2012). Specifically, female *Akt1*(-/-) mice exhibit increased hippocampal oscillation power in the theta, alpha, beta, and gamma frequency ranges (Chang et al. 2016). AKT1 also regulates GABAergic neuron differentiation and GABAAR expression, important for hippocampus-dependent cognitive functions (Chang et al. 2016). AKT1 interacts with S100B to promote neuronal cell proliferation (Arcuri et al. 2005). *S100b* knockout mice show a reduced γ band (30-80 Hz) response in the hippocampus after seizure induction with kainic acid (Sakatani et al. 2007). Subjects with medial temporal epilepsy show altered expressions of *S100B* (Lu et al. 2010). Overall, this is suggestive of some role for S100B-related pathways in the modulation of brain oscillations in specific conditions. *S100B* encodes a calcium-binding protein involved in neurite extension and axonal proliferation, ultimately being involved in synaptic plasticity and learning. Phosphorylation of Akt is enhanced by APOE (specifically, APOE3) (Okoro et al. 2016), whereas APOE interacts with S100B during astrocytic activation/inhibition ((Mori et al. 2005). *APOE* encodes a component of the Reelin signalling pathway, which has been involved in verbal memory deficits in SZ (Verbrugghe et al. 2012; Li et al. 2015). *APOE* is the most significant genetic risk factor for late-onset Alzheimer Disease (and thus, for the progressive decline in memory, executive function, and language, observed in this condition (Huynh et al. 2017). It has been suggested that *APOE* is also a candidate for primary progressive aphasia (see Rogalski et al. 2013 for discussion). Interestingly, *APOE* has been related to some of the metabolic changes that allowed bigger brains (and eventually, enhanced cognitive capacities) to emerge in our clade (Bufill and Carbonell, 2006). Interestingly too, the allele ε4 of the gene (consistently related to a higher risk for developing late onset Alzheimer’s disease) differentially affects low and high frequency bands (particularly, alpha, theta and delta) in several areas of the brain, plausibly accounting for the reduced cognitive abilities of the carriers (Canuet et al. 2012, Cuesta et al. 2015, Liang et al. 2017)

Signalling by NRG1 has been found to increase the expression of an isoform of DISC1, encoded by a robust SZ candidate, during neurodevelopment (Seshadriet al. 2010). DISC1, a protein containing multiple coiled coil motifs and located in the nucleus, cytoplasm and mitochondria, is involved in cortical development, callosal formation, and neurite outgrowth (Brandon and Sawa 2011; Osbun et al. 2011). *DISC1* has been associated to verbal reasoning in the general population (Thomson et al., 2014) and to category fluency in people with bipolar disorder (Palo et al., 2007) and schizophrenia (Nicodemus et al. 2014). Importantly, *DISC1* is regulated by FOXP2, a transcription factor encoded by the ‘language gene’ par excellence (Walker et al. 2012). θ-induced long-term potentiation is altered in the hippocampal area CA1 of transgenic mice expressing a truncated version of *Disc1* (Booth et al. 2014). Moreover, the inhibitory effect of DISC1 on NRG1-induced ERBB4 activation and signalling affects the interneuron-pyramidal neuron circuit (Seshadri et al. 2015).

Several other candidates for SZ are predicted to be functionally linked to DISC1 and/or NRG1, including CACNA1C, CACNA1I, COMT, and ZNF804A. All of these are known to impact oscillatory patterns. *CACNA1I* and *CACNA1C* encode different subunits of calcium channels. *CACNA1C* encodes the alpha 1C subunit of the Cav1.2 voltage-dependent L-type calcium channel, which contributes to β and γ wave generation (Kumar et al. 2015). Risk alleles of the gene correlate with lower performance scores in semantic verbal fluency tasks in schizophrenics (Krug et al. 2010). Pathogenic variants of *CACNA1C* have been identified in subjects with intellectual disability, executive dysfunction, hyperactivity-impulsivity that interferes with Attention-Deficit/Hyperactivity Disorder (ADHD) and/or ASD, as well as forms of childhood-onset epilepsy (Damaj et al. 2015). *CACNA1I* has been related to changes in sleep spindles in schizophrenics, a form of oscillation that constrains aspects of thalamocortical crosstalk, impacting on memory consolidation and learning (Manoach et al. 2015). Likewise, low voltage α has been associated with low activity levels in COMT, a catechol-O-methyltransferase that catalyzes the O-methylation of neurotransmitters like dopamine, epinephrine, and norepinephrine (Enoch et al. 2003). *COMT* has been regularly associated with language performance and processing, and language acquisition, particularly with verbal fluency (Krug et al. 2009, Soeiro-De-Souza et al. 2013, Sugiura et al. 2017), but also with reading abilities (Landi et al. 2013). Finally, ZNF804A, a zinc finger binding protein, modulates hippocampal γ oscillations and thus, the coordination of hippocampal and prefrontal distributed networks (Cousijn et al. 2015). It also contributes to cortical functioning and neural connectivity, because of its known role in growth cone function and neurite elongation (Hinna et al. 2015). SZ risk polymorphisms of *ZNF804A* result in lower performance scores in reading and spelling tasks (Becker et al. 2012), but also in task evaluating category fluency during semantic processing (Nicodemus et al. 2014). ASNP within intron 2 of the gene has been found to be associated with ASD subjects that are verbally deficient (Anitha et al. 2012).

Concerning *ERBB4*, this gene has been related to intellectual disability and speech delay (Kasnauskiene et al. 2013). ERBB4 is predicted to interact with PDGFRB, and putative homologs of these two genes have been found to interact in other species, particularly in *Drosophila melanogaster* and *Caenorhabditis elegans* (Figure 1). In human cells, a direct interaction of PDGFRB and one of the functional isoforms of ERBB4 has been recently documented (Sundwall et al 2010). *PDGFRB* encodes the β subunit of the platelet-derived growth factor (PDGF) receptor, which plays an important role in central nervous system development. In mice, the knockout of *Pdgfrb* results in reduced auditory phase-locked γ, which correlates with anatomical, physiological, and behavioural anomalies that are also found in schizophrenics, including decreased GABAergic compactness in the medial prefrontal cortex, the hippocampus, and the amygdala, deficient spatial memory and impaired social behaviour (Nguyen et al. 2011, Nakamura et al. 2015).

ERBB4 is also a functional partner of PTEN, a phosphatase that preferentially dephosphorylates phosphoinositide substrates. Both proteins collaborate in protrusion formation in rhombic lip cells (Sakakibara and Horwitz, 2006). Functional interactions are predicted as well between PTEN and PDGFRB (Figure 1). *PTEN* is a candidate for a subtype of ASD with macrocephaly which is usually present in conjunction with epilepsy (or paroxysmal EEG) (Buxbaum et al. 2007; Marchese et al. 2014). The gene is highlighted as a candidate for language deficits in ASD, because patients with PTEN-associated ASD show a delay in language development, characterised by poor processing speed and working memory (Naqvi et al. 2000, Tilot et al. 2015). PTEN is a major negative regulator of the mTOR signalling pathway, important for synaptic plasticity and neuronal cytoarchitecture (see Tilot et al. 2015 for review). The knockdown of *Pten* in mouse primary neuron cultures affects the expression of genes involved in neurogenesis, synaptic activity, and long-term potentiation (Lanz et al. 2013). In mice, the deletion of *Pten* in adult hippocampal neural stem cells increases proliferation and differentiation of stem cells toward hypertrophied neurons with abnormal polarity, causes seizures and macrocephaly, and impairs social behaviour (Amiri et al. 2012). Social dysfunction in mouse models of neural *Pten* loss includes repetitive behaviour, impaired emotional learning (in females) and increased anxiety (in males) (Page et al. 2009, Clipperton-Allen and Page 2014), but also seizures and epileptiform features (Ogawa et al. 2007). Interestingly, *Pten* deletion in mice ultimately yields deviant circuit formation in the dentate gyrus, responsible for excitation flow through the hippocampus (Pun et al. 2012), potentially impairing procedural memory capacities relevant to language.

PTEN is a strong partner of MAPK14, a p38 mitogen-activated protein kinase which is also functionally related to ERBB4 and PDGFRB (Figure 1). In glioma cells, the downregulation of *MAPK14* correlates with the upregulation of *PTEN*, resulting in the inhibition of cell migration in vitro (Dasari et al. 2010). The inhibition of MAPK14 activity suppresses hippocampal-dependent associative and spatial memory deficits in mouse models of synaptic dysfunction (Roy et al. 2015). Mice with a single copy disruption of *Mapk14* show protection against kainate-induced seizures (Namiki et al. 2007). Another partner of PTEN is SIRT1, a deacetylase of the sirtuin family, which negatively regulates neurogenesis and neural differentiation, contributes to axon formation and elongation, and plays a role in memory formation (Gao et al. 2010, Li et al. 2013, Saharan et al. 2013). Sirt1 is prevents seizures and seizure-induced damage in the hippocampus of rat models of epilepsy via miR activity (Wang et al. 2016). The gene is also highly expressed in the cochlea and the auditory cortex (Xiong et al. 2014). SIRT1 phosphorylation and activation by DYRK1A, a dual-specificity tyrosine phosphorylation-regulated kinase, promotes cell survival (Guo et al. 2010). *DYRK1A* is located within the Down Syndrome Critical Region within chromosome 21. In mice, *Dyrk1a* has proven to contribute to the balance between cortical and thalamic neurons (Guedj et al. 2012). *Dyrk1a* overexpression affects the expression of genes encoding GABAergic and glutamatergic related proteins, shifts the excitation/inhibition balance towards inhibition, and impacts on pathways involved in synaptogenesis and synaptic plasticity (Souchet et al. 2014), supporting a role of this gene in learning and memory (Hämmerle et al. 2003). *DYRK1A* has been related as well to lack of speech, mental retardation and microcephaly (Van Bon et al. 2011, Courcet et al. 2012). In mice, the upregulation of *Dyrk1a* also results in the upregulation of *Gad1* (Souchet et al. 2014), which encodes a glutamic acid decarboxylase that catalyzes the production of GABA, with a specific role in the development of GABAergic neurons in the hippocampus (Pleasure et al. 2000). *GAD1* has been related to the pathophysiology of SZ, but also to working memory deficits, because of its impact on prefrontal white matter structure (Lett et al. 2016). GAD1 is a target of FOXP2 (Konopka et al. 2009). Risk alleles of the gene impact as well on long-interval cortical inhibition (LICI) in the dorsolateral prefrontal cortex of schizophrenics, as showed by transcranial magnetic stimulation with electroencephalography (TMS-EEG): this suggests that the gene contributes to GABAergic inhibitory neurotransmission (Lett et al. 2016). Male *Gad1* (+/-) mice exhibit impaired social behavior (Sandhu et al., 2014).

GAD1 interacts with DLX5 and DLX6, two genes that encode homeobox transcription factors important for GABAergic interneuron development (Cobos et al. 2006; Ghanem et al. 2008; Poitras et al. 2010). Accordingly, in the developing ventral forebrain, the non-coding RNA Evf2 controls transcription of *Gad1*, *Dlx5*, and *Dlx6* through cis- and trans-acting mechanisms; *Evf2* mouse mutants exhibit reduced synaptic inhibition (Bond et al. 2009). *Dlx5* and *Foxp2* are expressed in the same intercalated cell masses of the amygdala in non-human primates and in rats, and in nearly the same neuronal populations of the striatum (Kaoru et al., 2010). *DLX5* and *DLX6* are core components of the gene network accounting for aspects of the evolution of our ability to learn and uses languages (see Boeck and Benítez-Burraco 2014 for review). Heterozygous mice for *Dlx5/6* exhibit reduced cognitive flexibility which appears to emerge from abnormal GABAergic interneurons and γ rhythms, particularly in fast-spiking interneurons (Cho et al. 2015), potentially contributing to the irregular long-lasting prefrontal and central γ in ASD, but also to SZ symptoms. Evf2 also recruits Mecp2 to DNA regulatory elements in the Dlx5/6 intergenic region (Bond et al. 2009), whereas *DLX5* has been reported to be modulated by MECP2 (Miyano et al. 2008). *MECP2* is the principal candidate for Rett syndrome, a neurodegenerative disease involving autistic behaviour, motor problems, and language loss (Uchino et al. 2001, Veenstra-VanderWeele and Cook 2004). MECP2 is a chromosomal protein that binds to methylated DNA and mediates transcriptional repression, and that is critically needed for normal function of GABA-releasing neurons (Chao et al. 2010). In mice, the loss of *Mecp2* from GABAergic interneurons results in auditory event-related potential deficits (Goffinet al. 2014). In response to auditory stimulation, *Mecp2*+/- mice recapitulate select γ and β band abnormalities and specific latency differences found in ASD subjects (Liao et al. 2012).

Another strong partner of PTEN (but also of MAPK14, PDGFRB, ERBB4, and NRG1) is EGR1, a transcription factor that contributes to neural plasticity and memory consolidation (Veyrac et al. 2014). *EGR1* is found induced in human epileptic foci and its expression levels correlate with the frequency, amplitude and area of the interictal spikes, a hallmark of epileptic neocortex (Rakhade et al. 2007). *EGR1* is a target of FOXP2 (Konopka et al. 2009). In turn, EGR1 downregulates *PLAUR* (Matsunoshita et al. 2011), which encodes the urokinase plasminogen activator receptor and which is also a target of FOXP2 (Roll et al. 2010). Mice lacking *Plaur* have significantly fewer neocortical GABAergic interneurons, which are vital for oscillatory processes (Bae et al. 2010), and exhibit nearly complete loss in parvalbumin-containing interneurons during brain development, which is associated with increased susceptibility to spontaneous seizures and with impaired social interactions (Bruneau and Szepetowski 2011). PLAUR is an effector of SRPX2, a protein with three sushi repeats motifs which is another of FOXP2 targets (Royer-Zemmour et al. 2008) and a candidate for rolandic epilepsy and speech dyspraxia (Roll et al. 2006). One distinctive feature of this benign type of epilepsy with an onset in childhood is the presence of abnormal centrotemporal sharp waves, an endophenotype of rolandic epilepsies that has been associated with *ELP4* (Strug et al. 2009). *ELP4* encodes one component of the elongator protein complex, involved in RNA transcription and tRNA modification, and important for cell mobility and migration, particularly during the development of the cerebral cortex (Creppe et al. 2009). Interestingly, the locus of *ELP4* has been linked to speech sound disorder (SSD) (Pal et al. 2010). Microdeletions of *ELP4* have also been associated with ASD and linguistic deficits (Addis et al. 2015).

*EGR1* expression is induced by CNR1 (Bouaboulaet al. 1995). Genomics studies have highlighted *CNR1* as an important gene for brain anomalies and metabolic changes in SZ (Yu et al. 2013, Suárez-Pinilla et al. 2015), for striatal response to happy faces in a Caucasian cohort of ASD people (Chakrabarti et al., 2006), and for cases of total absence of expressive speech (Poot et al. 2009). *CNR1* encodes the cannabinoid-1 receptor, which modulates θ and γ rhythms in different brain areas, such as the hippocampus, with an impact on sensory gating function in the limbic circuitry (Hajós et al. 2008). CNR1-positive GABAergic interneurons play an important role in the response to auditory cues, as well as in other aspects of behaviour (Brown et al. 2014). *CNR1* is functionally linked to several other genes encoding a subset of related proteins that also appears as a core component of our network (Figure 1), including *HTR1A*, *GRM8*, *GRIN2A*, *GRIN2B*, and *SHANK3*. Interestingly, most of these genes encode neurotransmitter receptors.

Beginning with *HTR1A*, this encodes the receptor 1A of serotonin and contributes to the modulation of hippocampal γ, influencing cognitive functions linked to serotonin, such as learning and memory (Johnston et al. 2014). Interestingly, the serotonin-1A receptor exhibits a lateralized distribution in the language areas, being the receptor binding 1.8-2.9% higher in right language areas and 2-3.6% higher in left auditory regions (Fink et al. 2009). *GRIN2A* and *GRIN2B* encode two components of the subunit NR2 of the NMDA receptor channel, involved in long-term potentiation, a physiological process underlying memory and learning. GRIN2A is reduced in fast-firing interneurons of schizophrenics, which contribute decisively to γ oscillation formation: a blockade of NR2A-containing receptors increases γ power and reduces the modulation of γ by low frequencies (Kocsis 2012). Language regression and speech impairments have also been found to result from *GRIN2A* mutations (Carvill et al. 2013, Lesca et al. 2013). The gene is also a candidate for rolandic epilepsies (Dimassi et al. 2014). Speech problems found in patients with mutations in *GRIN2A* include imprecise articulation, problems with pitch and prosody, and low performance on vowel duration and repetition of monosyllables and trisyllables, which are commonly diagnosed as dysarthria or dyspraxia (Turner et al. 2015). *GRIN2B* plays a key role in normal neuronal development and in learning and memory. Besides its involvement in SZ, ASD, and SLI, mutations in *GRIN2B* have been found in subjects with intellectual disability associated with behavioural problems and EEG anomalies, and in patients with epileptic encephalopathies which co-occur with impairment of motor and cognitive functions (Freunscht et al. 2013, Smigiel et al. 2016, Hu et al. 2016). Finally, *GRM8* encodes a protein with a glutamate, GABA-B-like receptor activity. Partial duplications of the gene have been associated to developmental delay and intellectual disability (DECIPHER patients 338209 and 289333). Several SNPs of the *GRM8* have been found associated with θ power in subjects with alcohol dependence, which suggests that variation in *GRM8* may modulate θ rhythms during information processing (Chen et al. 2009).

In several organisms, GRIN2B interacts with SHANK3, a postsynaptic scaffolding protein that seems to be important for the maintenance of the adequate balance between neuronal excitation and inhibition. Knockdown of *Shank3* in mouse primary neuron cultures affects the expression of genes involved in long-term potentiation and synaptic activity (Lanz et al. 2013). Cultured cortical neuron networks lacking *Shank3* show reduced excitation and inhibition behaviours (Lu et al. 2016). Specifically, mice lacking the exon 9 of the gene exhibit reduced excitatory transmission in the hippocampal CA1 region and increased frequency of spontaneous inhibitory synaptic events in pyramidal neurons, which result in mildly impaired spatial memory (Lee et al. 2015). Knocked out mice for the gene exhibit abnormal social interaction and repetitive grooming behaviour (Peça et al. 2011). SHANK3 has been linked as well to some of the distinctive symptoms of Phelan-McDermid syndrome (also known as 22q13 deletion syndrome), including intellectual disability, delayed or absent speech, autistic features, and seizures and abnormal EEG profiles (Soorya et al. 2013, Holder and Quach 2016).

Besides CNVs in *GRIN2A* and *SHANK3*, CNVs in genes related to SLI and DD have been found as well in patients with continuous spike and waves during slow-wave sleep syndrome and Landau-Kleffner syndrome, including *ATP13A4* and *CNTNAP2* (Lesca et al. 2012). The latter encodes a protein associated with K^+^ voltage-gated channels in pyramidal cells of the temporal cortex largely innervated by GABAergic interneurons (Inda et al. 2006). *CNTNAP2* additionally affects language development in non-pathological populations (see Whitehouse et al. 2011, Whalley et al. 2011, Kos et al. 2012). This effect is seemingly due to its role in dendritic arborization and spine development (Anderson et al. 2012), and in the regulation of cerebral morphology and brain connectivity (Scott-Van Zeeland et al. 2010, Tan et al. 2010, Dennis et al. 2011). Homozygous mutations or compound heterozygous CNVs of *CNTNAP2* are associated with speech and language regression and epilepsy (Strauss et al. 2006, Marchese et al. 2016, Smogavec et al. 2016). Interestingly, mice and rats with homozygous deletions of *Cntnap2* exhibit reduced spectral power in the α (9-12 Hz) range during wake (Thomas et al. 2016). In mice, *Cntnap2* is regulated by Auts2, a protein with a suggested role in cytoskeletal regulation (Oksenberg et al. 2014). *AUTS2* displays the strongest signal of a selective sweep in anatomically-modern humans compared to Neanderthals (Green et al. 2010, Oksenberg et al. 2013) and is a strong candidate for several neurodevelopmental disorders (see Oksenberg and Ahituv 2013 for review) Specifically, CNVs of the gene have been found in patients with language delay and seizures (Nagamani et al. 2013), and the gene has been cited as a candidate for epilepsy (Mefford et al. 2010). Interestingly, *AUTS2* has also been associated with differential processing speeds (Luciano et al. 2011).

As noted above, the dysfunction of GABA signalling contributes to ASD-like stereotypes, Rett syndrome phenotypes, and SZ (Chao et al. 2010, Fazzari et al. 2010). Abnormal changes in the GABA catabolism give rise to brain and behavioral disturbances that recapitulate the symptoms of ASD, including language impairment (Gibson et al. 1997, Pearl et al. 2003). The fact that our list of candidates for the language oscillogenome includes several receptors for GABA reinforces the view that GABA signalling is crucial for the oscillatory signature of language. As shown in Figure 1, a third subnetwork includes GABRB3, GABARAP and two interactors; NSF and SNAP25. *GABRB3* encodes the β-3 subunit of the GABA receptor A (Cook et al. 1998, Shao et al. 2002, Shao et al. 2003). Besides its known association with ASD, the gene has been associated as well with childhood absence epilepsy (Urak et al., 2006). Null mutations of *Gabrb3* in mice result in cleft palate and epilepsy (Homanics et al. 1997), whereas heterozygous deletion also encompassing Ube3a and Atp10a, recapitulates Angelman syndrome, a neurobehavioral disorder involving absence of language (Jiang et al. 2010). Differences in the expression level of the *GABRB3* have been related to changes in the firing of hippocampal pyramidal neurons and the activity of fast networks (Heisteket al. 2010). More generally, genetic variation in GABAA receptor properties have been linked to differences in β and γ oscillations, which seemingly impact on network dynamics and cognition (Porjeszet al. 2002). *GABARAP* is a candidate for dyslexia and encodes a GABAA receptor-associated protein involved in the clustering of neurotransmitter receptors, but also in inhibitory neural transmission. *Gabarap* knockout mice exhibit abnormal paroxysmal sharp waves in the hippocampus (Nakajima et al. 2012). Estrogen depletion resulting from the inhibition of the dyslexia candidate *CYP19A1*, a member of the cytochrome P450 family that catalyzes the formation of aromatic C18 estrogens from C19 androgens, affects GABA synthesis and gives rise to increased spine density and decreased threshold for hippocampal seizures (Zhou et al. 2007). Regarding NSF and SNAP25 (the former a candidate for dyslexia and the latter a candidate for SZ, ASD, and dyslexia), both are needed for neurotransmitter release and synaptic function. *NSF* encodes a protein involved in vesicle-mediated transport in the Golgi apparatus, whereas SNAP25 contributes to the formation of the soluble NSF attachment protein receptor complex. In mice, reduced levels of Snap25 seems to be related to more frequent spikes, diffuse network hyperexcitability, and epileptiform discharges, as well as to cognitive deficits and social impairment (Corradini et al. 2014, Braida et al. 2015).

Downregulation of GABA receptors has been linked as well to altered expression of *FMR1*. Specifically, reduced levels of GABRβ3 and of FMRP have been found in the vermis of adult subjects with ASD (Fatemi et al. 2011), as well as in the hippocampus of *En2*(-/-) mice model of ASD (Provenzano et al. 2015). FMRP, a polyribosome-associated RNA-binding protein, is encoded by *FMRP1*, the main candidate for Fragile X syndrome, a condition involving language deficits and frequent features of ASD (Kaufmann et al. 2004, Smith et al. 2012). Low levels of FMRP have been found as well in schizophrenic patients with low IQs (Kovács et al. 2013). *Fmr1-*knockout mice exhibit enhanced mGluR5 signalling, which results in altered neocortical rhythmic activity because of changes in neocortical excitatory circuitry (Hays et al. 2011). These mice also exhibit abnormal patterns of coupling between θ and γ oscillations in perisomatic and dendritic hippocampal CA1 local field potentials, resulting in abnormally weak changes during tasks involving cognitive challenge (Radwan et al. 2016). Also, inhibitory dysfunctions in layer II/III of the somatosensory cortex has been found in *Fmr1* knockout mice, in particular, a reduced activation of low-threshold-spiking interneurons and reductions in synchronized synaptic inhibition and coordinated spike synchrony in pyramidal neurons in response to mGluR agonists (Paluszkiewiczet al. 2011).

*FMR1* has been suggested to fit with *ROBO1*, *KIAA0319*, *S100B*, and *DCDC2*, among others, into a theoretical gene network important for neurite outgrowth and neuronal migration (Poelmans et al. 2011). All these genes are candidates for DD according to results from association studies, GWA analyses, and CNVs studies (Paracchini et al. 2016), and all have been related to abnormal patterns of brain oscillations or seizures when mutated. Accordingly, they seem to us to be promising candidates for the oscillatory signature of language. Rare variants in the intergenic region between *DCDC2* and *KIAA0319*, and in one intron of *DCDC2*, which encodes a doublecortin domain-containing protein (*locus* DYX2) have been associated with differences between dyslexic and control children in a late mismatch negativity around 300-700 ms originating in right central-parietal areas when discriminating between complex auditory stimuli, such as syllables and words (Czamara et al. 2011). The protein encoded by *ROBO1* is a membrane receptor of the immunoglobulin superfamily which contributes to regulate interaural interaction in auditory pathways (Lamminmäki et al. 2012). *ROBO1* is targeted by miR-218, which is found downregulated in the hippocampus of people suffering from medial temporal lobe epilepsy (Kaalund et al. 2014).

The remainder of our candidate genes are not clearly functionally interconnected in the core interacting network (Figure 1), although all of them play relevant roles in brain oscillations and are candidates for the basis of language impairments (see Table 1). This is why we still regard them as important components of the language oscillogenome. *FOXP1*, which encodes a forkhead box transcription factor, is co-expressed with *FOXP2* in some areas of the brain and the protein FOXP1 forms heterodimers with the FOXP2 protein. *FOXP1* haplo-insufficiency has been found in patients with epileptiform discharges, severe speech delay, and delayed gross motor skills (Carr et al. 2010). *Cx3cr1* knockout mice show deficient synaptic pruning, weak synaptic transmission, decreased functional brain connectivity, and social and behavioural features that resemble those found in ASD patients (Zhan et al. 2014). Interestingly, these mice also exhibit reduced θ-driven connections between prefrontal cortex and dorsolateral hippocampus relative to wild-type littermates (Zhan 2015). *CX3CR1* encodes a receptor for fractalkine, a chemokine and transmembrane protein important for cell adhesion and migration. Specifically, CX3CR1 has been involved in the crosstalk between the microglia and the neural stem/progenitor cells during adult hippocampal neurogenesis, which is important for memory, learning and cognition (de Miranda et al. 2017), to the extent that CX3CR1 deficiency results in impairment of hippocampal cognitive function and synaptic plasticity (Rogers et al. 2011). As with other proteins associated with ion channels, like CNTNAP2, DPP10 is of interest due to its binding capacity to K^+^ channels and its ability to modify their expression and biophysical properties (Djurovic et al. 2010). Rare mutations in *DPP10* have been associated with ASD (Marshall et al. 2008). Interestingly, the transcription start site of the gene is hypermethylated in the neurons of the prefrontal cortex of humans compared to extant primates, and regulatory sequences at *DPP10* carry elevated nucleotide substitution rates and regulatory motifs absent in archaic hominins, with signals of recent selective pressures and adaptive fixations in modern populations (Shulha et al. 2012). This reinforces the view that this gene might have contributed to cognitive abilities and disorders that are unique to humans *MBD5* encodes a protein with a methyl-CpG-binding domain which binds to methylated DNA. *MBD5* haplo-insufficiency has been associated with epilepsy, severe speech delay, mental retardation, and ASD-features (Williams et al. 2010, Talkowski et al. 2011). The gene encodes a methyl-CpG-binding protein. *SETBP1*, which encodes a SET binding protein, is a candidate for Schinzel-Giedion syndrome, which entails severe developmental delay and occasional epilepsy (Ko et al. 2013, Miyake et al. 2015). Mutations on the gene also result in social and behavioural problems (Coe et al. 2014). As a candidate for SLI, GWAs studies have associated *SETBP1* to the complexity of linguistic output (Kornilov et al. 2016). Microdeletions causing the disruption of the gene impact mostly on expressive abilities, whereas receptive language is quite preserved, to the extent that some patients are able to communicate using gestures and mimics (Filges et al. 2011, Marseglia et al. 2012). The C-terminal portion of COL4A2 arrests cell proliferation and migration; mutations in DD-candidate *COL4A2* have been found in patients suffering from epilepsy and severe developmental delay (Giorgio et al. 2015, Smigiel et al. 2016). *SCN1A* encodes the large α subunit of the voltage-gated sodium channel NaV1.1, which plays a key role in the generation and propagation of action potentials. Mutations in *SCN1A* have been found in people with ASD (Weiss et al. 2003, O’Roak et al. 2011), but mostly in patients with epilepsy (Schutte et al. 2016). The gene is also associated with Dravet syndrome, a condition characterised by cerebellar signs and a deficit in expressive language with relatively spared comprehension, resulting from motor speech problems which affect to motor planning and executing (Battaglia et al. 2013, Turner et al. 2017). In mice, the downregulation of *Scn1a* disturbs hippocampal oscillations and impairs spatial memory (Bender et al. 2013). Finally, *KANSL1*, which encodes a putative transcriptional regulator involved in the acetylation of nucleosomal histone H4, plays a role in gene transcription regulation and chromatin organization as part of the NSL1 complex. *KANSL1* is a candidate for Koolen-de Vrries syndrome, which entails epilepsy and developmental delay with moderate intellectual disability, which impacts mostly on expressive language abilities (Koolen et al. 2016).

The functional enrichment, based on gene ontology (GO) annotations performed using String algorithm, of our set of candidates for the language oscillogenome (Table 2) points out that most of these genes act in signalling pathways found to be of significance for language processing via its oscillatory implementation, particularly through dopaminergic, GABAergic and glutamatergic synapses (see Boeckx and Benítez-Burraco 2014a, b for discussion). The top-scoring biological processes (resulting from functional annotations) include the regulation of cognitive process of particular relevance for language, including learning and memory. Lastly, regarding the cellular localization of the proteins, the majority of them seem to be found in the plasma membrane, inside the neuron projection components, confirming their role as regulators of neuronal interconnection. In the next section, we discuss how the role played by these genes may underlie most of the oscillatory aspects of brain function that are important for language production and comprehension.

**Table 2.** Functional enrichment of the entire set of candidates for the oscillogenome. according to Gene Ontology (GO) consortium annotations. This enrichment was performed using String algorithm for gene network analysis. A false-discovery rate cutoff of 0.05, obtained after Bonferroni correction, was set to select significant functions. Except for the “molecular function” annotation, only the top fifteen scoring categories are displayed.

| **Biological Process (GO)** |  |  |  |
| --- | --- | --- | --- |
| *pathway ID* | *pathway description* | *count in gene set* | *false discovery rate* |
| GO:0044708 | single-organism behavior | 16 | 2.07e-12 |
| GO:0007610 | behavior | 16 | 5.66e-11 |
| GO:0030182 | neuron differentiation | 19 | 2.25e-10 |
| GO:0048468 | cell development | 22 | 9.17e-10 |
| GO:0048666 | neuron development | 17 | 1.16e-09 |
| GO:0007611 | learning or memory | 11 | 2.03e-09 |
| GO:0007613 | memory | 9 | 2.03e-09 |
| GO:0007399 | nervous system development | 23 | 2.69e-09 |
| GO:0023051 | regulation of signaling | 26 | 2.69e-09 |
| GO:0008284 | positive regulation of cell proliferation | 16 | 6.28e-09 |
| GO:0031175 | neuron projection development | 15 | 6.28e-09 |
| GO:0050804 | modulation of synaptic transmission | 11 | 6.28e-09 |
| GO:0048167 | regulation of synaptic plasticity | 9 | 8.92e-09 |
| GO:0030030 | cell projection organization | 17 | 8.99e-09 |
| GO:0048699 | generation of neurons | 19 | 8.99e-09 |
| **Molecular Function (GO)** |  |  |  |
| *pathway ID* | *pathway description* | *count in gene set* | *false discovery rate* |
| GO:0048156 | tau protein binding | 3 | 0.000993 |
| GO:0019904 | protein domain specific binding | 8 | 0.00284 |
| GO:0005102 | receptor binding | 11 | 0.0299 |
| GO:0008066 | glutamate receptor activity | 3 | 0.0351 |
| **Cellular Component (GO)** |  |  |  |
| *pathway ID* | *pathway description* | *count in gene set* | *false discovery rate* |
| GO:0098589 | membrane region | 18 | 2.79e-09 |
| GO:0098590 | plasma membrane region | 17 | 2.79e-09 |
| GO:0097458 | neuron part | 18 | 3.09e-09 |
| GO:0098805 | whole membrane | 22 | 3.09e-09 |
| GO:0043005 | neuron projection | 16 | 5.2e-09 |
| GO:0044463 | cell projection part | 16 | 5.2e-09 |
| GO:0042995 | cell projection | 20 | 1.46e-08 |
| GO:0097060 | synaptic membrane | 10 | 4.91e-08 |
| GO:0036477 | somatodendritic compartment | 13 | 9.06e-08 |
| GO:0044297 | cell body | 10 | 9.05e-06 |
| GO:0030424 | axon | 9 | 1.43e-05 |
| GO:0045202 | synapse | 11 | 1.43e-05 |
| GO:0034702 | ion channel complex | 8 | 1.52e-05 |
| GO:0044456 | synapse part | 10 | 1.85e-05 |
| GO:0030425 | dendrite | 9 | 3.65e-05 |

**Table 2. Functional enrichment of the entire set of candidates for the oscillogenome.**
| **KEGG Pathways** |  |  |  |
| --- | --- | --- | --- |
| *pathway ID* | *pathway description* | *count in gene set* | *false discovery rate* |
| 04728 | Dopaminergic synapse | 7 | 4.36e-06 |
| 04727 | GABAergic synapse | 5 | 0.000209 |
| 04015 | Rap1 signaling pathway | 6 | 0.000513 |
| 04068 | FoxO signaling pathway | 5 | 0.000513 |
| 04724 | Glutamatergic synapse | 5 | 0.000513 |
| 05031 | Amphetamine addiction | 4 | 0.000778 |
| 04080 | Neuroactive ligand-receptor interaction | 6 | 0.00155 |
| 04020 | Calcium signaling pathway | 5 | 0.00187 |
| 04713 | Circadian entrainment | 4 | 0.00187 |
| 04723 | Retrograde endocannabinoid signaling | 4 | 0.00217 |
| 05033 | Nicotine addiction | 3 | 0.00272 |
| 05014 | Amyotrophic lateral sclerosis (ALS) | 3 | 0.00526 |
| 04010 | MAPK signaling pathway | 5 | 0.00657 |
| 05214 | Glioma | 3 | 0.00813 |
| 04720 | Long-term potentiation | 3 | 0.00834 |

In other to check the language-specificity of our network, we compared the functional annotations of our network with the result of the functional enrichment analyses of the whole set of candidates for the four conditions also considered in our work (SZ, ASD, DD and SLI). Because the String algorithm did not provide with significant results for DD and SLI, we instead relied on Enrichr (http://amp.pharm.mssm.edu/Enrichr/). The results are shown in table 3.

**Table 3.**
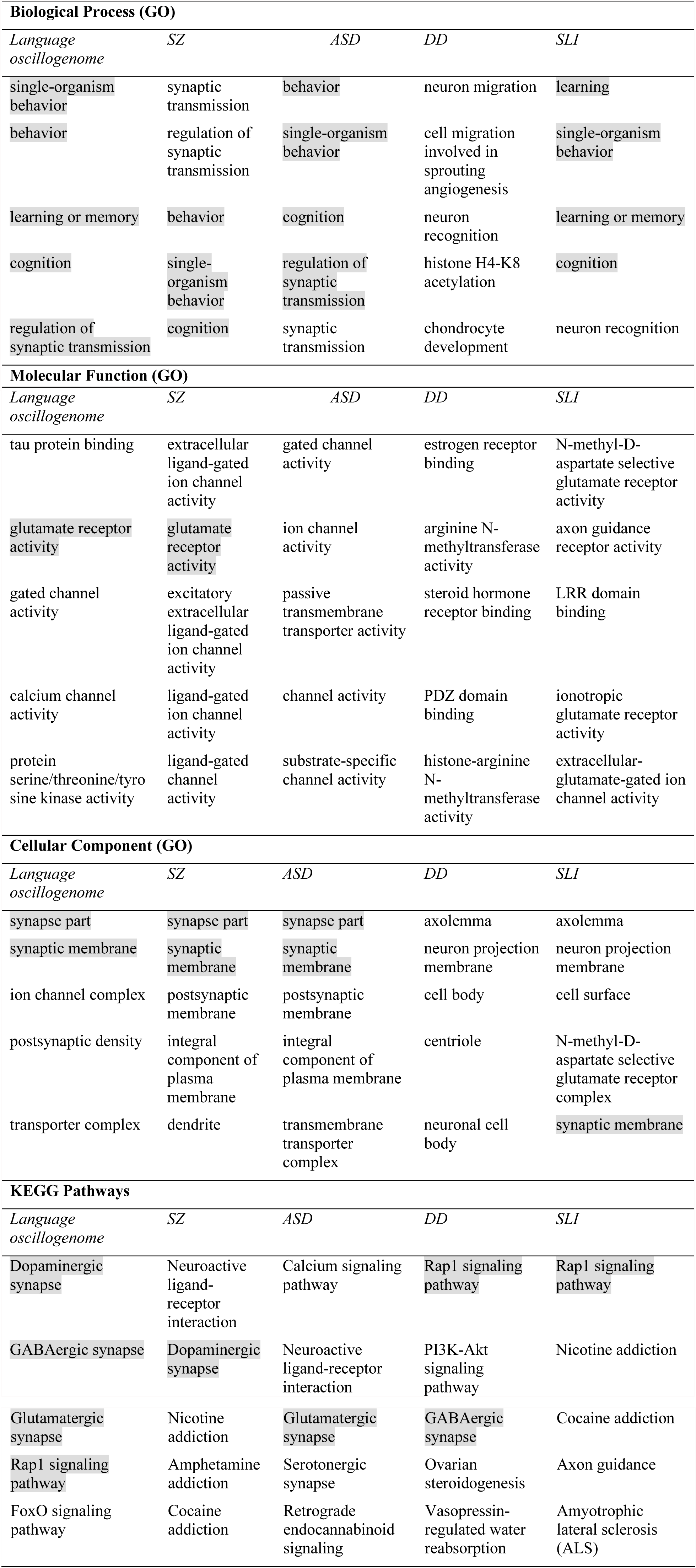
Comparison of the functional enrichment of candidates for the oscillogenome. SZ, ASD, DD, and SLI. This enrichment was performed using the Enrichr algorithm for gene network analysis. A false-discovery rate cutoff of 0.05, obtained after Fisher exact test correction, was set to select significant functions. Only the top five scoring categories are displayed. Categories are ranked according to their adjusted p value, which derives from running the Fisher exact test for many random gene sets in order to compute a mean rank and standard deviation from the expected rank for each term in the gene-set library and finally calculating a z-score to assess the deviation from the expected rank.

| **Biological Process (GO)** |  |  |  |  |
| --- | --- | --- | --- | --- |
| *Language oscillogenome* | *SZ* | *ASD* | *DD* | *SLI* |
| single-organism behavior | synaptic transmission | behavior | neuron migration | learning |
| behavior | regulation of synaptic transmission | single-organism behavior | cell migration involved in sprouting angiogenesis | single-organism behavior |
| learning or memory | behavior | cognition | neuron recognition | learning or memory |
| cognition | single-organism behavior | regulation of synaptic transmission | histone H4-K8 acetylation | cognition |
| regulation of synaptic transmission | cognition | synaptic transmission | chondrocyte development | neuron recognition |
| **Molecular Function (GO)** |  |  |  |  |

| *Language*<br>*oscullogenome* | *SZ* | *ASD* | *DD* | *SLI* |
| --- | --- | --- | --- | --- |
| tau protein binding | extracellular<br>ligand-gated<br>ion channel<br>activity | gated channel<br>activity | estrogen receptor<br>binding | N-methyl-D-aspartate selective<br>glutamate receptor<br>activity |
| glutamate receptor<br>activity | glutamate<br>receptor<br>activity | ion channel<br>activity | arginine N-methyltransferase<br>activity | axon guidance<br>receptor activity |
| gated channel<br>activity | excitatory<br>extracellular<br>ligand-gated<br>ion channel<br>activity | passive<br>transmembrane<br>transporter activity | steroid hormone<br>receptor binding | LRR domain<br>binding |
| calcium channel<br>activity | ligand-gated<br>ion channel<br>activity | channel activity | PDZ domain<br>binding | ionotropic<br>glutamate receptor<br>activity |
| protein<br>serine/threonine/tyro<br>sine kinase activity | ligand-gated<br>channel<br>activity | substrate-specific<br>channel activity | histone-arginine<br>N-methyltransferase<br>activity | extracellular-<br>glutamate-gated ion<br>channel activity |

Cellular Component (GO)
| *Language*<br>*oscullogenome* | *SZ* | *ASD* | *DD* | *SLI* |
| --- | --- | --- | --- | --- |
| synapse part | synapse part | synapse part | axolemma | axolemma |
| synaptic membrane | synaptic<br>membrane | synaptic<br>membrane | neuron projection<br>membrane | neuron projection<br>membrane |
| ion channel complex | postsynaptic<br>membrane | postsynaptic<br>membrane | cell body | cell surface |
| postsynaptic density | integral<br>component of<br>plasma<br>membrane | integral<br>component of<br>plasma membrane | centriole | N-methyl-D-aspartate selective<br>glutamate receptor<br>complex |
| transporter complex | dendrite | transmembrane<br>transporter<br>complex | neuronal cell<br>body | synaptic membrane |

**Table 3. Comparison of the functional enrichment of candidates for the oscillogenome.**
| *Language*<br>*oscullogenome* | *SZ* | *ASD* | *DD* | *SLI* |
| --- | --- | --- | --- | --- |
| Dopaminergic<br>synapse | Neuroactive<br>ligand-receptor<br>interaction | Calcium signaling<br>pathway | Rap1 signaling<br>pathway | Rap1 signaling<br>pathway |
| GABAergic synapse | Dopaminergic<br>synapse | Neuroactive<br>ligand-receptor<br>interaction | PI3K-Akt<br>signaling<br>pathway | Nicotine addiction |
| Glutamatergic synapse | Nicotine addiction | Glutamatergic synapse | GABAergic synapse | Cocaine addiction |
| Rap1 signaling pathway | Amphetamine addiction | Serotonergic synapse | Ovarian steroidogenesis | Axon guidance |
| FoxO signaling pathway | Cocaine addiction | Retrograde endocannabinoid signaling | Vasopressin-regulated water reabsorption | Amyotrophic lateral sclerosis (ALS) |

As expected, the top-scoring biological processes resulting from the GO analyses are quite similar across groups, supporting the view that although learning and memory are key aspects of language acquisition and processing, their impairment can result in different cognitive disorders. Regarding the cellular components in which our candidates for the language oscillogenome are preferably found, they are specifically involved in ion channel formation and function, contrary to candidates for SZ, ASD, DD, or SLI, reinforcing the view that our candidates are important for aspects of brain rhythmicity. That said, we believe that the language-specificity of our oscillogenomic candidates might rely on the pathways to which they contribute. Specifically, they are components of the dopaminergic signalling pathway, which is usually highlighted as important for motor behaviour and vocal learning (Kameda et al. 2011). More importantly, they contribute as well to GABAergic signalling, which is relevant for the maintenance of our species-specific cognitive profile (Long et al. 2013). As discussed in detail in Boeckx and Benítez-Burraco (2014a), some of the key changes that contributed to the emergence of our ability to learn and use languages (usually referred to as our language-readiness) involved GABAergic signalling. Among them we wish highlight the evolutionary changes in the core candidate for globularization of the human skull/brain, namely, *RUNX2*, also involved in the development of hippocampal GABAergic neurons (Pleasure et al., 2000), and the generation of an extra source of GABAergic neurons resulting from the expansion of dorsal thalamic structures (Striedter 2004, Rakic 2009, Geschwind and Rakic 2013). Importantly, as reasoned in Murphy and Benítez-Burraco (2016b), hippocampal θ waves, which are produced by slow pulses of GABAergic inhibition (Vertes and Kocsis 1997), are involved in the extraction of language feature-sets from memory. Additionally, we expect that the language-specificity of our network rely as well on the role played by our candidates in FOXO and RAP1 pathways, contrary to the genes related to broader cognitive conditions like SZ and ASD. Regarding the FOXO signalling pathway, we have already discussed the relevance of *FOXP1* and *FOXP2*. We wish also highlight *FOXO1*, which is a target of the two core candidates for the evolutionary changes that prompted the emergence of our language-readiness, namely, *RUNX2* (Kuhlwilm et al. 2013) and *FOXP2* (Vernes et al. 2011) (see Boeckx and Benítez-Burraco 2014a, 2014b, and Benítez-Burraco and Boeckx 2015 for details). Additionally, FOXO1 upregulates *RELN* (Daly et al. 2004), a candidate for ASD and for lissencephaly with language loss (Hong et al. 2000, Wang et al. 2014), and is phosphorylated by DYRK1A (Huang and Tindall, 2007). Regarding the RAP1 signalling pathway, it is important for regulating MAPK activity and for promoting GABA(B) receptor surface expression (Zhang et al. 2015). As shown in Table 3, candidates for clinical conditions affecting language only are enriched in components of this pathway.

In order to delve into the language-specificity of our network, we also used Enrichr to generate the expression grids of our candidates for the oscillogenome across the brain regions profiled by the Allen Brain Atlas (http://www.brain-map.org/). Figure 2 compares the grids for up- and downregulated genes in the brain, with the grids for SZ, ASD, DD, and SLI. Overall, our candidates are most upregulated in the medial septal nucleus, which innervates the hippocampal formation and which plays a key role in the generation of θ waves (Pignatelli et al. 2012). They are also highly upregulated in the thalamus (specifically, in the sensory-motor cortex, but also in the stria terminalis, which serves as a major output pathway of the amygdala), the insula (in the ventral and the dorsal parts of the agranular insular area and in the superficial stratum of insular cortex), and in the striatum (in the septopallidal area, the striatal septum and the pallidal septum). They are significantly downregulated in several parts of the cerebellum, including the cerebellar cortex and the vermis. On the contrary, candidates for SLI are mostly upregulated in the basal forebrain and the hindbrain, whereas they are most downregulated in different areas of the cortex, including the retrosplenial, frontal, and parietal cortices. We think that this might contribute to confer an oscillomic specificity to our language-related set of candidates compared to other candidates for language dysfunction. As we discuss in detail in Boeckx and Benítez-Burraco (2014a), the thalamus may have played a central role in the evolutionary changes resulting in our language-readiness. The thalamus acts as a relay device for most of the brain areas involved in language processing, particularly, between the cortex and the basal ganglia (Lieberman 2002, Theyel et al. 2009), and it is crucially involved in aspects of the syntax-semantics interface (Wahl et al. 2008; David et al. 2011). Importantly for our hypothesis, the thalamus controls the oscillations generated in the cortex (Theyel et al. 2009, Saalmann et al., 2012). Specifically, thalamic input allows for an enrichment of oscillatory activity in different frequency bands (Singer 2013, Uhlhaas et al. 2013, Cannon et al. 2014), and it is thalamic GABAergic neurons that crucially contribute to restrict cortical activity, by providing low-frequency oscillations that are capable of embedding higher-frequency oscillations across distant brain areas (Whitman et al. 2013). This embedding plays a key role in aspects of language processing (see Boeckx 2013a,b and Murphy 2015b, 2016b for details). Regarding the striatum, it is crucially involved in vocal learning, as part of the cortico-thalamic-striatal circuits responsible for motor planning and procedural learning (for details, see Boeckx and Benítez-Burraco 2014b).

**Figure 2.**
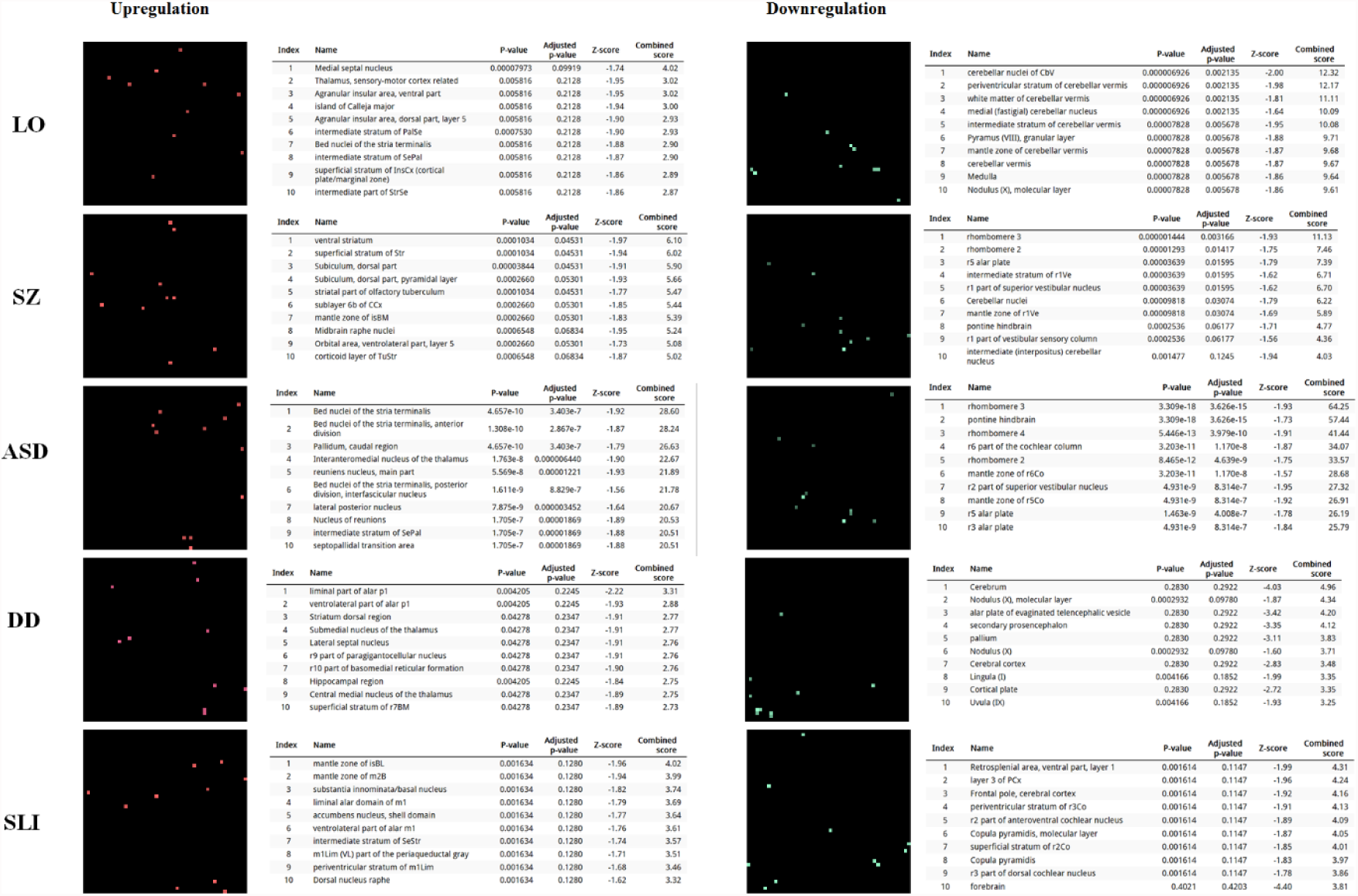
Expression profiles in the brain of the set of genes considered in this study. Expression grids were generated with Enrichr. Brain regions where genes are most upregulated are displayed in red, whereas region in which genes are most downregulated are shown in green (the more light the colour, the more up- or downregulated a gene is). The tables contain the most up- or downregulated genes according to the Combined Score. The acronyms for the brain regions are from the Allen Brain Atlas and can be checked at http://developingmouse.brain-map.org/docs/Legend_2010_03.pdf.

We have equally examined the brain expression profiles of our candidates across development, in order to know whether they are down- or upregulated during the time window when changes in the brain wiring and function, important for language acquisition, take place. As shown in figure 3, some of these genes exhibit expression peaks in different brain areas between 0-5 years of age, like *APOE*, *CX3CR1*, *CYP19A1*, and *PDGFRB*. Conversely, others are significantly downregulated after birth, like *COL4A2*, *DLX5*, *GRIN2B*, *ROBO1*, or *SETBP1*. Interestingly, several of our candidates exhibit dissimilar expression trends in different brain areas. For instance, after birth, *CACNA1L* is upregulated in the striatum and downregulated in the thalamus. Likewise, *CX3CR1* is downregulated in the cerebellum and upregulated in the cortex, the thalamus, and the striatum. Because of the involvement of the thalamus in our model of language processing and language evolution, it is also of interest the expression profile of *CYP19A1* in this area, with a neat peak during childhood. Also of interest is *HTR1A*, with shows consecutive expression peaks across development: it is highly expressed in the embryonic cortex and the embryonic hippocampus (with a second peak during adolescence), it is upregulated in the cerebellum around birth, and it is upregulated in the amygdala in young people. We expect that these patterns of expression contribute to explain the observed changes in brain activity during language processing across developmental stages, and ultimately, account for some of the changes in the language abilities of the child.

**Figure 3.**
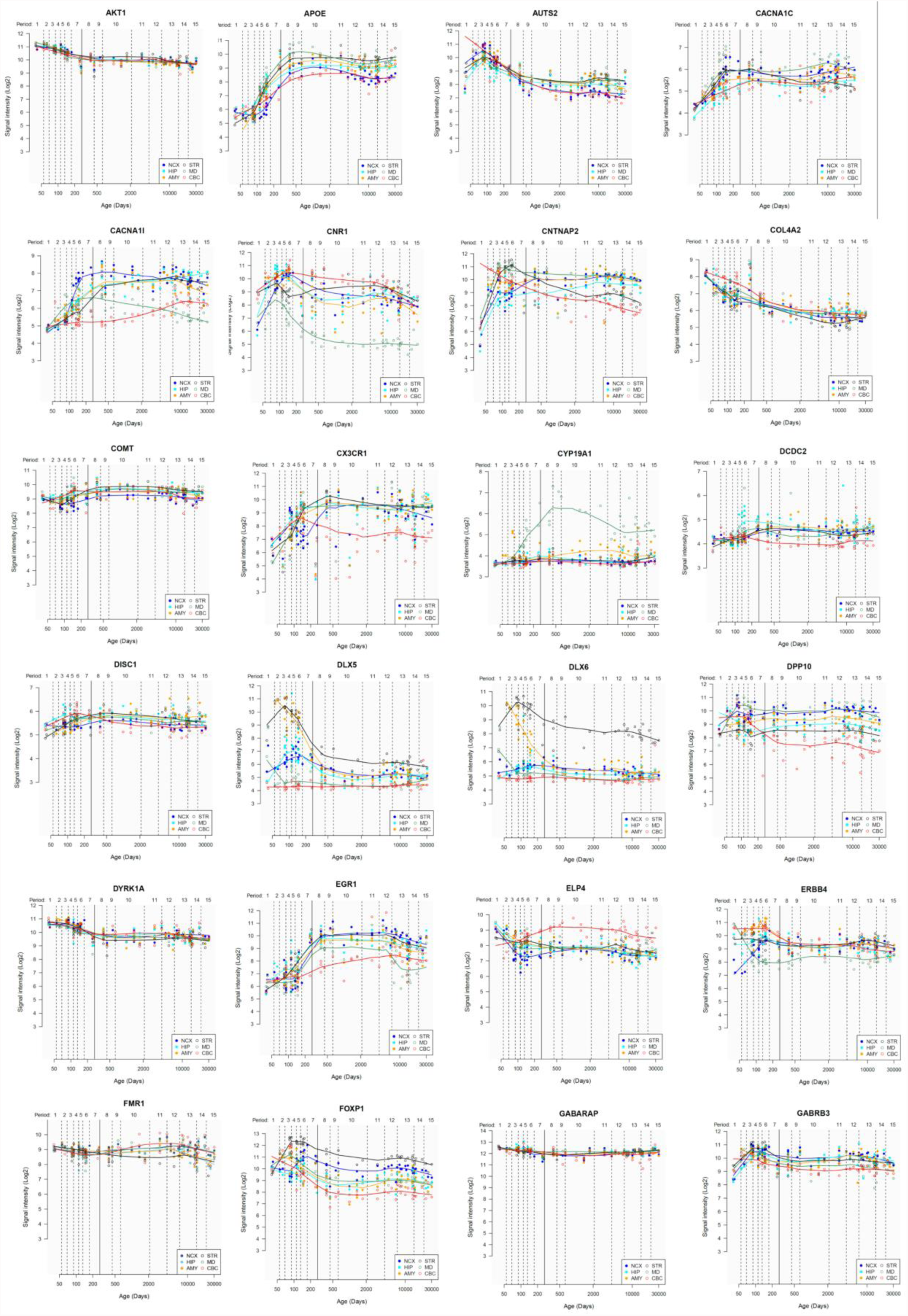
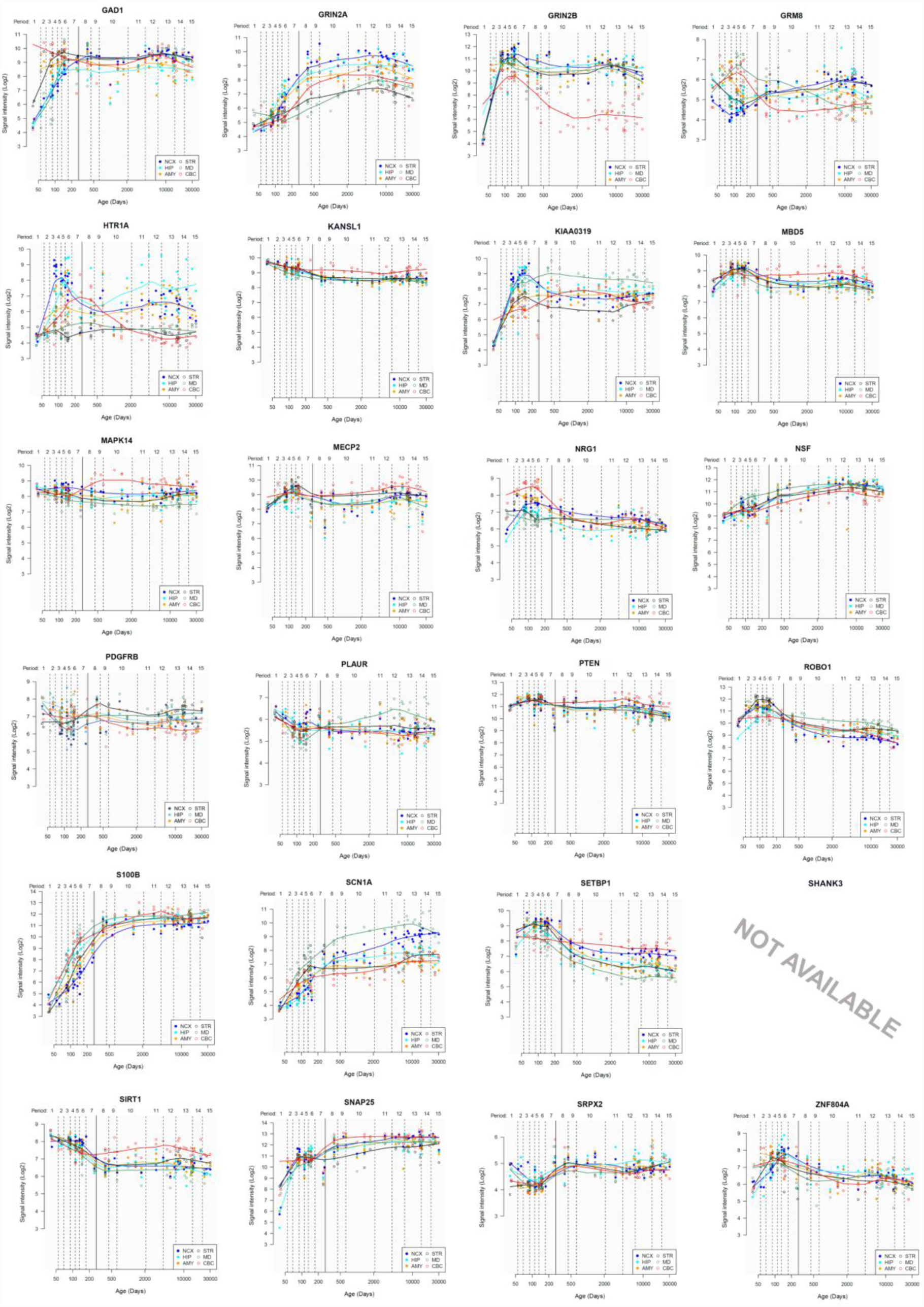
Individual brain expression profiles across development of the set of genes considered in this study. The expression data are from the Human Brain Transcriptome Database (http://hbatlas.org/). Six different brain regions are considered: the cerebellar cortex (CBC), the mediodorsal nucleus of the thalamus (MD), the striatum (STR), the amygdala (AMY), the hippocampus (HIP) and 11 areas of neocortex (NCX). Data for *SHANK3* were not available.

## 3. Linking the language oscillogenome to language processing

Having documented the most likely candidates which could constitute a robust, testable oscillogenome, we now turn to the neurocomputational implementation of language processing, and how an abnormal genetic profile can in turn give rise to abnormal oscillatory signatures. The core feature of our oscillogenomic approach is a rich level of cross-disciplinary integration.

As Anderson (2016: 6) says of the relationship between evolutionary psychology and neuroscience, ‘function in the brain depends upon, at least: a neural network, an underlying genetic network, and an overlaid chemical gradient. Each of these elements is only partially understood, and their dynamic interactions even less so’. By attempting to draw causal relations between genes, oscillations, and linguistic computations we hope that we can shed some light on the nature of these interactions. As noted earlier, the interpretation and construction of linguistic phrases require a range of particular cross-frequency couplings across certain regions (Murphy 2016a, b). Genes are expected to contribute decisively to the emergence of this global neuronal workspace, yielding specific patterns of long-distance connections among distributed neurons and, as a result, specific oscillatory signatures of language.

In Murphy (2016b), an ‘oscillomic’ model was proposed through which linguistic representations are itemized via γ nesting within parahippocampal θ cycles. This complex would then be nested within the phase of left-cortical δ, attributing to the set a phrasal identity. Certain of these γ clusters would then slow to β to be maintained in memory. This process of phrasal construction is assumed in Murphy (2015b) to be the only human-specific linguistic computation. Ewerdwalbesloh et al. (2016) show that holding visually constructed objects in memory (as opposed to ‘whole’, presented object) results in greater fronto-parietal θ synchronisation. Since the maintenance of constructed objects in fronto-parietal circuits is vital for language, it was hypothesized in Murphy (2016b) that the language system might recruit this (possibly generic) neural code for computationally analogous purposes. In brief, a lexicalisation process generated by a θ-γ code would interact with a phrasal construction process of δ phase-entrainment. This is a more computationally explicit framework than predictive coding models (e.g. Kesller et al 2016), going beyond simple procedures like ‘what’ and ‘where’ computations into set-theoretic notions more in line with contemporary linguistic theory.

What are the reasons to believe that oscillations have any causal-explanatory power with respect to language? Vosskuhl et al. (2015) used transcranial alternating current stimulation (tACS) to decrease participant’s θ such that the θ:γ ratio was altered and a larger number of γ cycles could be nested within θ. This carefully controlled study demonstrated that this manipulation increased working memory performance. Although this study does not directly speak to syntactic combinatorics, given the reliance the language system has on working memory we can infer that the θ-γ code is causally related to linguistic representation construction.

Other experimental evidence for the role of this neural code in language comes from a number of places, such as the finding that left-cortical δ entrains to hierarchical linguistic structures from syllables to sentences (Ding et al. 2016). In addition, the roles ascribed here to particular oscillations and oscillatory interactions are supported by the broader (often domain-general) roles argued for by Ketz et al. (2015); namely, that θ is implicated in recollective memory, β in executive control, and α in sensory information gating. In addition, α decreases at right fronto-temporal sites occur when syllables are temporally expected (Wilsch et al. 2015). Such expectancy effects appear in other linguistic domains, with increased semantic predictability leading to reduced parieto-occipital α (Wöstmann et al. 2015). Finally, a verbal generation task by Wojtecki et al. (2016) resulted in 6-12Hz power increases and enhanced θ-α coherence between the subthalamic nucleus and frontal sites as a function of successful task performance. Despite the relative paucity of experimental work, the role of α in semantic and phonological prediction seems clear – in particular given this rhythm’s function in coordinating the representations constructed by θ-γ coupling (as reviewed in Murphy 2016b).

Language impairments in ASD at the syntax-semantics interface most often involve difficulties with relative clauses, wh-questions, raising and passives (Perovic and Janke 2013, Perovic et al. 2007, Wada 2015). Increased γ power has also been found for individuals with autism (Kikuchi et al. 2013; Rojas et al. 2008), likely going some way to explain their abnormal linguistic comprehension given the model discussed in Murphy (2016b), with Kikuchi et al. finding in addition reduced cross-cortical θ, α and β. Bangel et al. (2014) discovered lower β power during a number estimation task in individuals with ASD, and broader rhythmic abnormalities have been found. These findings may be (partly) explained through what we have reviewed above; namely, that low voltage α has been associated with low activity levels in COMT (Enoch et al. 2003). As mentioned, *ZNF804A*, *HTR1A* and *GRIN2B* modulate hippocampal γ oscillations (Cousijn et al. 2015) with *ZNF804A* additionally contributing to cortical functioning and neural connectivity, and so these genes may play a role in the etiopathogenesis of the ASD γ profile. We also noted that knockout of *Pdgfrb* results in reduced auditory phase-locked γ oscillations, which may be a primary cause of similar oscillatory effects in ASD and SZ. We also reviewed how θ-induced long-term potentiation is altered in hippocampal area CA1 of transgenic mice expressing a truncated version of *Disc1* (Booth et al. 2014).

The ASD oscillome also appears to frequently involve reduced θ during tasks necessitating inter-regional synchronisation (Doesburg et al. 2013). The reduced θ in the ASD population (also documented by Kikuchi et al. 2013) may therefore arise from these or related hippocampal ensembles, which would in turn contribute to working memory deficits, impacting semantic processing.

Abnormally long-lasting prefrontal and central γ is exhibited by individuals with ASD during processing semantic incongruities (Braeutigam et al. 2008), which potentially reflects the execution of a general search mechanism (high γ) to replace the normal rhythmic processes (low γ) to extract and compare representations. As noted, heterozygous mice for *Dlx5/6* exhibit reduced cognitive flexibility which appears to emerge from abnormal GABAergic interneurons and γ rhythms (Cho et al. 2015), and it is possible that this is the correct oscillogenomic model to account for this abnormal γ profile.

ASD patients with abnormal levels of *MECP2* show an abnormal γ response in auditory stimulus discrimination tasks (Peters et al. 2015). Similarly, in response to auditory stimulation mice with a heterozygous loss of *Mecp2* function display increased latency of cortically sourced components, and also display γ and β abnormalities associated with ASD and SZ (Liao et al. 2012). Picture-naming tasks also lead to lower left inferior frontal γ and β power in ASD subjects relative to neurotypical controls (Buard et al. 2013), and rhythmic connectivity between auditory and language cortices is also abnormal (Jochaut et al. 2015); results potentially explicable via this oscillogenomic account. In particular, Jochaut et al. discovered that speech processing results in severely impaired θ-γ coupling in ASD (Jochaut et al. 2015), a finding which may relate to the knockout of *Pdgfrb* resulting in reduced auditory phaselocked γ. In addition, we noted that *Fmr1* knockout mice exhibit enhanced mGluR5 signalling, resulting in altered neocortical rhythmic activity (Hays et al. 2011). Since these mice exhibit abnormal patterns of coupling between hippocampal θ and γ (Radwan et al. 2016), this provides another strong oscillogenomic candidate for θ-γ coupling disruptions.

A study of lexical decision in SZ also exposed lower left-temporal and left-frontal α and β power (Xu et al.’s 2013) – a rhythmic profile also found in Moran and Hong (2011) and Uhlhaas et al. (2008). A sentence presentation task by Xu et al. (2012) revealed reduced θ at occipital and right frontal lobe sites. As noted, the cannabinoid-1 receptor encoded by *CNR1* modulates θ and γ rhythms in several brain areas (Hajós et al. 2008) and so may be involved in these abnormalities. Relatedly, a blockade of NR2A-containing receptors increases γ power and reduces low-frequency γ modulation; we have previously documented unusually fast γ in SZ and ASD patients (Murphy and Benítez-Burraco 2016), and so this may be part of the underling oscillogenomic basis. Decomposing the P300 event-related component into its constituent θ and δ rhythms, Jones et al. (2004) report significant linkage and linkage disequilibrium between frontal θ band and a single nucleotide polymorphism from the cholinergic muscarinic receptor gene (*CHRM2*) on chromosome 7. Due to the likely role of this gene in higher cognition (Gosso et al. 2007), this makes it a strong candidate gene for cognitive deficits in SZ.

Knockout of *Ppargc1a* in mice decreases the spread of activation in hippocampal CA1 and limits pyramidal cell spiking, giving rise also to unusual modulations of kainate-induced γ oscillations (Bartley et al. 2015). *PPARGC1A* deficiency in ASD may consequently lead to direct oscillatory alterations at this frequency band. We also noted an association between *GRM8* and θ power, suggesting that variations in *GRM8* may modulate θ rhythms during information processing, potentially opening it up as a candidate gene for ASD, SZ and DD, given the abnormal θ modulations documented in these disorders.

With respect to the oscillatory basis of linguistic prosody, we noted that speech problems found in patients with mutations in *GRIN2A* include imprecise articulation and problems with pitch and prosody – archetypal problems documented in DD. Other research indicates that individuals with developmental DD cannot achieve correct phonological representations in the brain, and that these problems arise from impaired phase-locking to slower modulations in the speech signal (below 10Hz, particularly around 2Hz), impacting syllabic parsing (Hämäläinen et al. 2012, see also Lehongre et al. 2011). Due to its relevance in the P300 component, the cholinergic muscarinic receptor gene *CHRM2* is a possible candidate for these δ abnormalities (Callaway 1983). Soltész et al. (2013) observed weaker entrainment in right auditory cortex of dyslexic patients during the processing of tone streams delivered at 2 Hz. The authors suggested a connection between reading performance and anticipatory δ phase-synchronization. Abnormal δ rhythms in auditory cortex have been found in dyslexics during the processing of speech sounds too (Molinaro et al. 2016).

It has also been suggested that increased anterior β is strongly reflective of dysphonetic dyslexics (with grapheme-to-phoneme conversion difficulties) whereas increased posterior β are typically found in dyseidetic children (with problems accessing the visual lexicon) (Flynn et al. 1992). These findings are compatible with the model proposed in Murphy (2016b) and discussed above, since anterior β is here assumed to be involved in the maintenance of the existing ‘cognitive set’, with abnormal β impairing the ability of dysphonetic dyslexics to hold one linguistic representation in memory and compare/convert it into another.

Relative to neurotypicals, dyslexics additionally display stronger high γ phase-synchronization in left auditory cortex, possibly indicating the wealth of spectrotemporal information reaching this region, compromising the θ-related auditory buffering capacity along with verbal working memory (Lehongre et al. 2011). This would in turn impair the feature-set combinatorial capacities of dyslexics, with both θ and γ, and their cross-frequency coupling, being abnormal, and so the potential candidate genes discussed above for ASD and SZ (e.g. *GRM8*) are also possible candidates for dyslexia.

Turning to SLI, Bishop et al. (2010) explored the discrimination of non-linguistic sounds in a group of 32 patients, comparing them to syllables in an oddball paradigm. Healthy controls exhibited event-related desynchronization in δ, θ and α during the presentation of oddballs, but SLI patients did not, pointing to a low-level auditory perceptual impairment in SLI. Further studies are needed in order to develop a more fine-grained picture of the perceptual and computational properties of SLI language comprehension, but we can nevertheless conclude that the candidate genes discussed above for these frequency bands remain potential candidates for the SLI oscillogenome.

Comparing the language deficits observed in dyslexia and SLI on the one hand with those observed in ASD and SZ on the other, it seems clear that they both exhibit δ abnormalities of distinct neurocomputational properties. In dyslexia, entrainment to the speech envelope (phonology) is impaired due to abnormal δ (leading to problems with slow-rate speech processing; Goswami et al. 2013), whereas in ASD and schizophrenia it appears that patients cannot properly exploit the δ-related processes of phrasal construction and identification. The δ rhythm therefore seems to act as a syntax-phonology interface but also a syntax-semantics interface, depending on which other regions are impaired in the particular disorder, such as the right supramarginal gyrus (Cutini et al. 2016) and left inferior frontal gyrus (Molinaro et al. 2016) in dyslexics.

Because many of the the computational roles we have ascribed to oscillations are notably generic (with the exception of language-specific δ-driven phrasal construction), this genericity of neurocognitive function can potentially map on to the Generalist Genes Hypothesis (Plomin & Kovas 2005; Kovas & Plomin 2006), such that the genes we have here claimed to be relevant to specific linguistic deficits (be it syntactic, phonological, and so on) are also likely implicated in other cognitive capacities, by virtue of the generic computational roles of, for instance, the θ-γ code or α inhibition.

There are alternative accounts to the one presented here which downplay (or simply reject) the existence of any causal-explanatory role for oscillations in language. For instance, Goucha et al. (2017) claim that ‘those [oscillatory] mechanisms seem to already be in place in other species. For example, despite the crucial brain expansion that took place in primates and especially humans compared to other mammals, the rhythmical hierarchy of oscillations is mainly kept unchanged’. While it may be true that this hierarchy has ‘mainly’ been kept unchanged – as indeed recent neuroethological work by Kikuchi et al. (2017) attempts to show – thus far there have been no neuroethological experiments using the kind of stimuli which would allow researchers to compare the oscillatory basis of putatively human-specific computations (like phrase-structure building) to the neural responses of non-human primates attempting to interpret or parse identical structures. Kikuchi et al. simply expose humans and monkeys to artificial grammars; not only does this not guarantee that the human subjects would recruit their language systems, but even if it did Kikuchi et al.’s data analysis did not investigate the kind of cross-frequency couplings over the particular regions and rhythms claimed here to be responsible for phrase-structure building. For instance, Kikuchi et al. only examine coupling between low frequencies and γ (see their Figure 4), and not coupling between low frequencies such as δ, θ and β (or indeed phase-phase coupling, claimed in Murphy 2016b to potentially play a role in syntactic processing). The authors found that ‘learned ordering relationships modulate the observed form of neural oscillatory coupling in [humans and monkeys]’, but this is a far cry from interpreting or generating hierarchically structured expressions. Indeed, further testing the model outlined here could involve not only refining Kikuchi et al.’s study to involve a broader class of stimuli and analyses, but it could also involve using MEG and EEG to investigate the oscillatory responses in people with autism and schizophrenia during language comprehension, going some way to lend more direct, experimental forms of support.

As a way of modelling what we have discussed here, Figure 4 outlines a general schema with some specific examples taken from this section. The bridge between the three levels described here – between the pathophysiology of language deficits in particular cognitive disorders and their linguistic profiles – remains very much open, but we hope that our framework will play a reflective role in theoretically grounding emerging findings in genetics and neural dynamics within a broader understanding of language processing and evolution.

**Figure 4.**
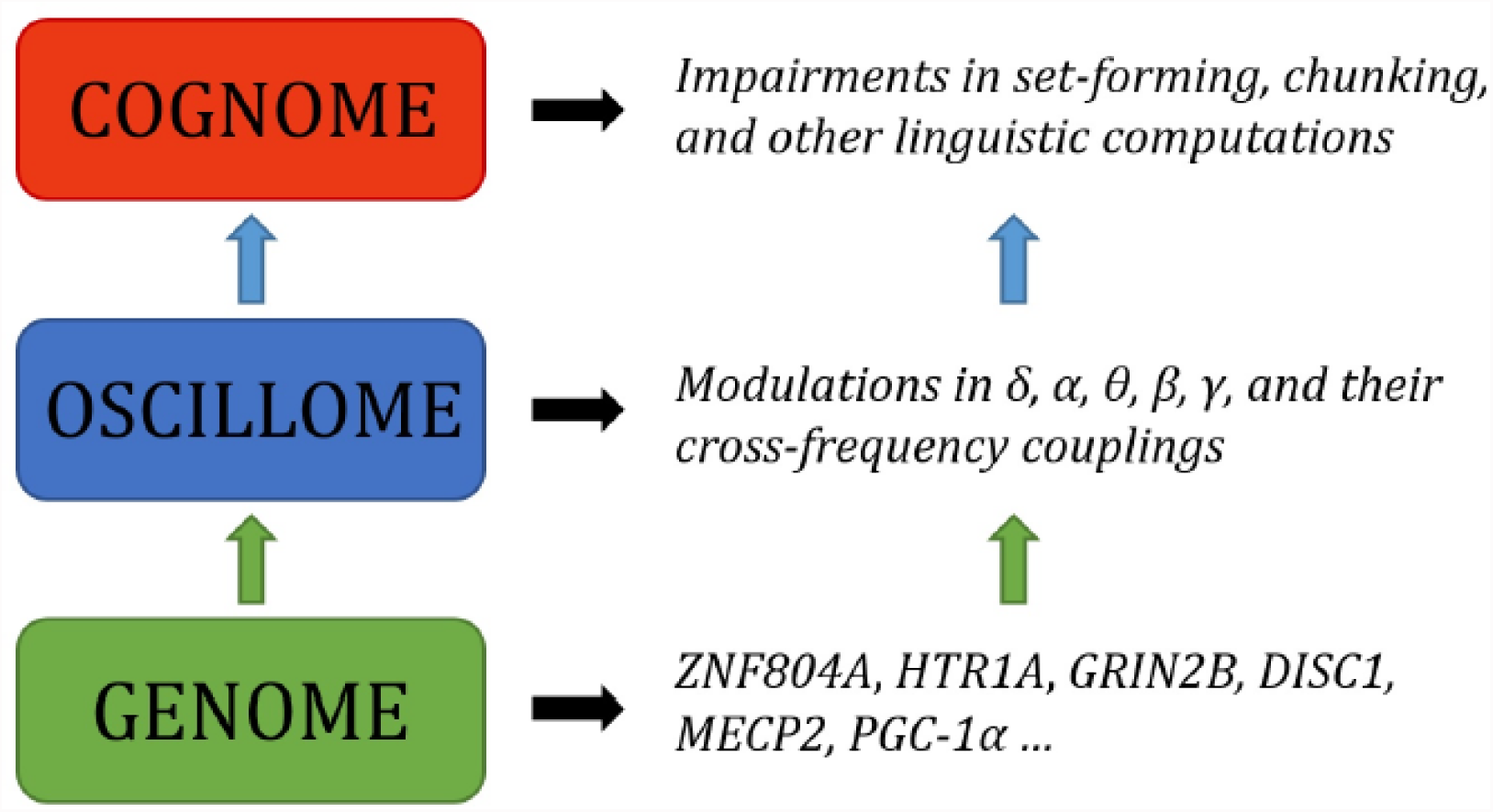
Outline of a putative oscillogenomic model for the human faculty of language. ASD, SZ, SLI and DD have been used as guiding ‘oscillopathies’, whereas linguistic theory has been employed as a guide for the neurocomputational basis of language. The genome is expected to modulate frequency bands and their interactions at the level of the oscillome, which in turn impacts computational operations at the ‘cognome’, that is, the basic cognitive operations underlying language, to use a term of Poeppel’s (2012). For instance, as noted in the text, knockout of *Ppargc1a* in mice decreases the spread of activation in hippocampal CA1 and limits pyramidal cell spiking, leading to unusual modulations of kainate-induced γ. *PPARGC1A* deficiency in ASD may consequently lead to direct oscillatory alterations at this frequency band; a hypothesis pending experimental confirmation.

## 4. Conclusions

Contemporary sequencing technologies have greatly expanded the set of genes associated with cognitive conditions entailing language deficits. The polygenism seen in these diseases is somewhat commensurable with the polygenism expected for language, which is necessary to properly characterize if we want to understand how language unfolds in the brain and develops in the child. Molecular biology techniques have significantly increased our knowledge of the role played by these genes in the healthy and the impaired brains. Neuroimaging facilities show how the brain organizes through development and processes language, in effect, via the coupling of diverse brain oscillations which enables complex interactions between local and distant brain areas. Nonetheless, it remains imperative to bridge the gap between genes, brain development and function, and language; and consequently to bridge the gap between pathological mutations, abnormal brain activity, and language deficits (Figure 5). Because brain oscillations can provide an explanation for how the brain processes language, but can also construct successful endophentypes of conditions involving language impairment, we have here relied on them to attempt to make substantial theoretical gains in this direction. There are also a number of potentially fruitful avenues to follow with respect to testing some of the core claims made here. For instance, if a given population (e.g. individuals with dyslexia) are known to exhibit particular genetic disruptions which are relevant to the generation of certain brain rhythms (e.g. δ), then this population could undergo M/EEG testing to determine if the linguistic capacities discussed here (e.g. phonological processing) are indeed disrupted in the manner predicted.

**Figure 5.**
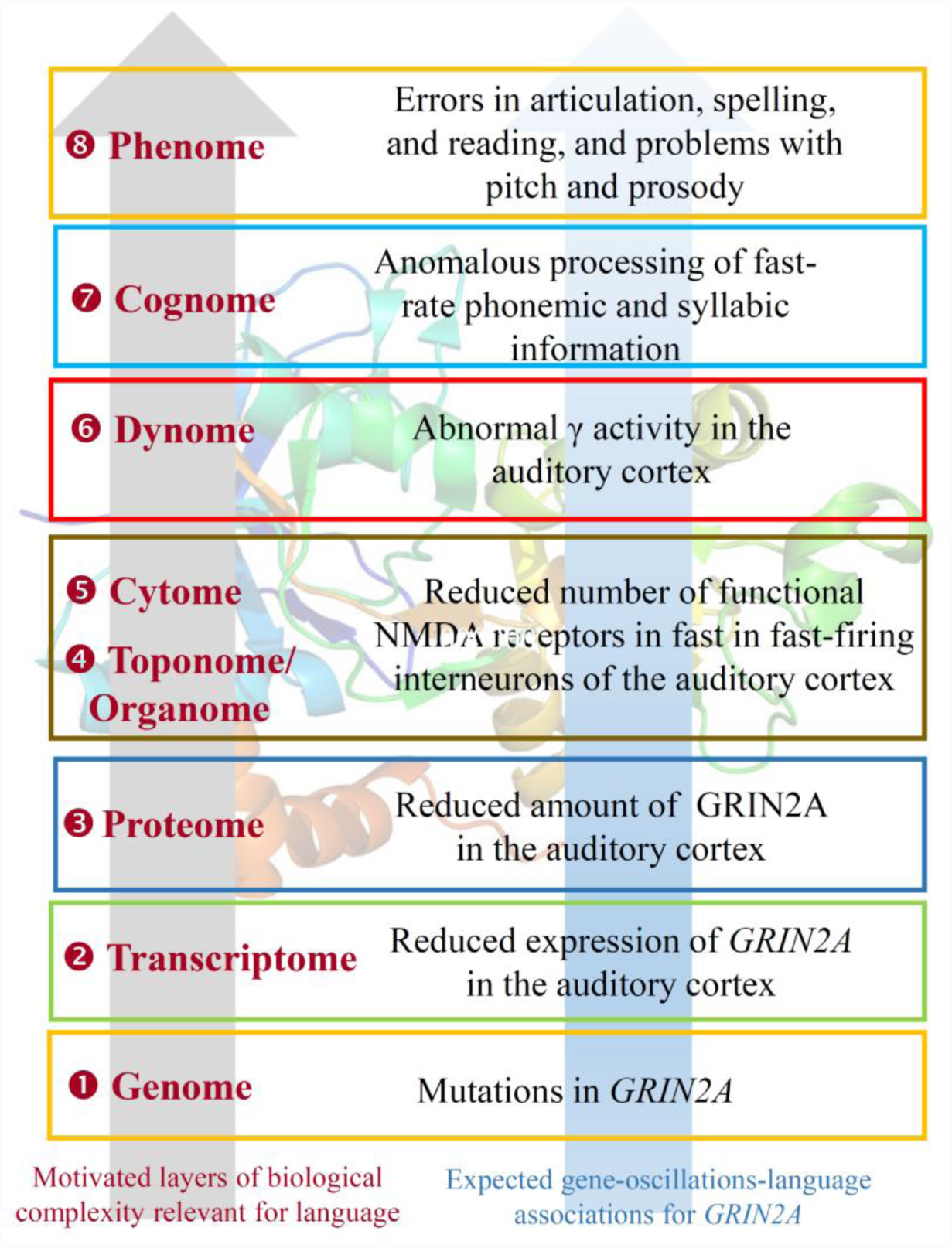
*GRIN2A* as an example of the observed (and expected) bridging links across the different levels of the biological analysis of language. As highlighted in the text, mutations in the gene give rise to different forms of epilepsy-aphasias, like Landau-Kleffner syndrome, continuous spike and waves during slow-wave sleep syndrome, and rolandic epilepsies. *GRIN2A* expression is found in several brain areas; a noteworthy upregulation of the gene is observed during late embryonic development. *GRIN2A* encodes the subunit 2A of the NMDA receptor, which plays a key role in long-term potentiation, a physiological process important for memory and learning. This role seemingly results in part from its effect on γ oscillation formation and modulation. As also noted in the text, GRIN2A levels are reduced in fast-firing interneurons of people with schizophrenia. As also discussed, mutations in *GRIN2A* result in errors in articulation, and in problems with pitch and prosody, which pertain to linguistic prosody, and which can be tracked to the abnormal γ activity, which is crucial for the correct processing of fast-rate phonemic and syllabic information. The 3D structure of GRIN2A, inserted as the background of the picture, is from the RSCB Protein Data Bank (http://www.rcsb.org/pdb/home/home.do). Here, ‘genome’ refers to the set of genes related to brain rhythms that are relevant for language, ‘transcriptome’, to their RNA products, and ‘proteome’ to the proteins they encode. ‘Toponome’ refers to the whole set of codes of proteins and other biomolecules found in the cells surface, whereas ‘Organome’ refers to the set of cell signalling molecules involved in cell and organ crosstalk. ‘Cytome’ refers to the collection of different cell types of the organism. ‘Connectome’ refers to the wiring of brain areas involved in language processing. ‘Dynome’ refers to the brain dynamics underlying (and supporting) this processing, in the line of Kopell et al., 2014 and Murphy, 2015. ‘Cognome’ refers to the basic cognitive operations underlying language (and in this case, speech processing), in the line of Poeppel, 2012. Finally, ‘phenome’ refers to the discrete, language-specific activities (in this case, phonological and phonetic aspects of speech).

Our main conclusion is that the functions of the genes discussed here crucially match aspects of the language oscillome. We have argued that the molecular findings appear to align with the experimental oscillatory results, which in turn align with components of the language cognitive phenotype. As it stands, these findings only afford tenuous causal-explanatory power to the present genome-oscillome-language linking hypotheses, and further experimental oscillatory and genetic research is required to strengthen the viability of the current gene set and increase the number of candidate genes. Specifically, we need to refine the mapping of these and other future candidate genes on specific cell functions, brain areas, aspects of brain function, neuronal developmental processes, and basic cognitive abilities. Animal models will help refine our understanding of the role of these genes and reinforce the links we have highlighted in the paper. Lastly, we expect that our approach will help gain a better understanding of the complex etiopathogenesis of cognitive conditions entailing problems with language, which should help in turn to design better therapeutic approaches to the diseases (see Wilkinson and Murphy 2016) aimed to ameliorate the symptoms and improve the abilities of the affected people.

## Acknowledgments

This work was supported by an Economic and Social Research Council scholarship (1474910) to EM, and by funds from the Spanish Ministry of Economy and Competitiveness (grant numbers FFI2014-61888-EXP and FFI-2013-43823-P) to ABB.

